# Single-cell multiome and enhancer connectome of human retinal pigment epithelium and choroid nominate pathogenic variants in age-related macular degeneration

**DOI:** 10.1101/2025.03.21.644670

**Authors:** Sean K. Wang, Jiaying Li, Surag Nair, Reshma Korasaju, Yun Chen, Yuanyuan Zhang, Anshul Kundaje, Yuwen Liu, Ningli Wang, Howard Y. Chang

## Abstract

Age-related macular degeneration (AMD) is a leading cause of vision loss worldwide. Genome-wide association studies (GWAS) of AMD have identified dozens of risk loci that may house disease targets. However, variants at these loci are largely noncoding, making it difficult to assess their function and whether they are causal. Here, we present a single-cell gene expression and chromatin accessibility atlas of human retinal pigment epithelium (RPE) and choroid to systematically analyze both coding and noncoding variants implicated in AMD. We employ HiChIP and Activity-by-Contact modeling to map enhancers in these tissues and predict cell and gene targets of risk variants. We further perform allele-specific self-transcribing active regulatory region sequencing (STARR-seq) to functionally test variant activity in RPE cells, including in the context of complement activation. Our work nominates new pathogenic variants and mechanisms in AMD and offers a rich and accessible resource for studying diseases of the RPE and choroid.

## INTRODUCTION

Age-related macular degeneration (AMD) is a leading cause of vision loss affecting an estimated 200 million people worldwide.^1^ AMD can be divided into two types: non-exudative (“dry”), characterized by the formation of extracellular deposits called drusen and progressive atrophy of the retinal pigment epithelium (RPE) and photoreceptors; and exudative (“wet”), characterized by pathologic neovascularization of the choroid. Decades of research have led to effective anti-angiogenic therapies for wet AMD.^2–5^ However, treatment options for dry AMD remain limited, and the underlying disease mechanisms are still poorly understood.

Twin studies estimate the heritability of AMD to range from 46-71%,^6,7^ suggesting that genetic analyses of AMD such as genome-wide association studies (GWAS) should reveal disease targets. Indeed, a landmark GWAS by Fritsche et al. in 2016 with over 30,000 subjects identified 52 genomic variants independently associated with AMD, including several encoding amino acid changes in complement factors.^8^ Nonetheless, most variants uncovered by Fritsche et al. are of unknown functional significance, largely because they reside in noncoding regions of the genome. For these variants, it can be unclear which cell types they affect, which genes they regulate, and whether they are truly causal or rather linked to a nearby variant driving the association.

To decipher the role of noncoding variants, we and others have adopted a multiomic framework combining techniques such as single-cell RNA sequencing (scRNA-seq) and single-cell assay for transposase-accessible chromatin sequencing (scATAC-seq).^9–11^ By using scRNA-seq to define cell types and integrating scATAC-seq to map chromatin accessibility, this approach can pinpoint the specific cellular contexts in which noncoding variants might function. In parallel, methods characterizing cis-regulatory elements have been harnessed to locate variants with possible gene regulatory activity. Studies of the 3D genome that detect physical enhancer-gene connections, such as HiChIP for histone H3 lysine 27 acetylation (H3K27ac),^12^ have been used to predict interactions between variants and their target genes.^13,14^ The impact of variants on transcription has also been experimentally tested using massively parallel reporter assays (MPRAs) and self-transcribing active regulatory region sequencing (STARR-seq),^15^ which employ high-throughput measurement of enhancer activity to nominate causal variants.^16,17^

Here, we applied a multiomic framework to the RPE and choroid, the earliest sites of AMD pathology,^18–20^ to systematically analyze nearly 2,000 coding and noncoding variants implicated in this disease. Using joint scRNA-seq and scATAC-seq, we generated a single-cell gene expression and chromatin accessibility atlas of human RPE and choroid from eyes with and without AMD. We then performed H3K27ac HiChIP and Activity-by-Contact (ABC) modeling^21^ to map enhancers in these tissues and conducted allele-specific STARR-seq to functionally test the regulatory potential of variants in RPE cells. Our work proposes new pathogenic variants and mechanisms in AMD and provides a rich and accessible resource for studying diseases of the RPE and choroid.

## RESULTS

### Single-cell multiomics reveal gene expression and chromatin accessibility landscapes of RPE and choroid cell types

To generate a single-cell multiome of human RPE and choroid, we performed joint scRNA- and scATAC-seq profiling of RPE-choroid complexes from 14 postmortem eyes of nine individuals (Figure 1A; Table S1). Four eyes from three individuals had a diagnosis of dry AMD, and none of the eyes had any ocular disease other than dry AMD or cataract. Nuclei capture from RPE and choroid was initially hampered by excess pigment interfering with single-cell microfluidics, which may explain why prior single-cell studies of these tissues featured relatively few cells.^22–24^ To overcome this limitation, we modified the Omni-ATAC protocol^25^ to optimize separation of pigment from RPE and choroid nuclei by incorporating a second density gradient and more stringent filtration, enabling much higher cell yields (Protocol S1). Following quality control filtering (Figures S1A-F) and doublet removal (Figure S2A), we resolved 12 clusters, including three corresponding to cell types from the ciliary epithelium and neural retina anatomically adjacent to the RPE and choroid (Figure S2B).^26–28^ Upon excluding these three clusters, we obtained a total of 39,326 human RPE and choroid cells in ten clusters which we assigned to nine distinct cell types (Figure 1B; Figure S2C). RPE cells and fibroblasts were the most abundant cell types detected, while B cells were the rarest, constituting only 0.7% of profiled cells (Figure 1C; Table S2). For each cell type, we identified specific marker genes consistent with those from published scRNA-seq data of human RPE and choroid (Figure S3A; Data S1).^22^ These included *RPE65* for RPE cells, *FBLN1* for fibroblasts, *TYR* for melanocytes, *FLT1* for vascular endothelial cells, *F13A1* for macrophages, *CD247* for T cells, *SCN7A* for Schwann cells, *ACTA2* for pericytes and smooth muscle cells, and *BANK1* for B cells (Figure 1D).

**Figure 1.**
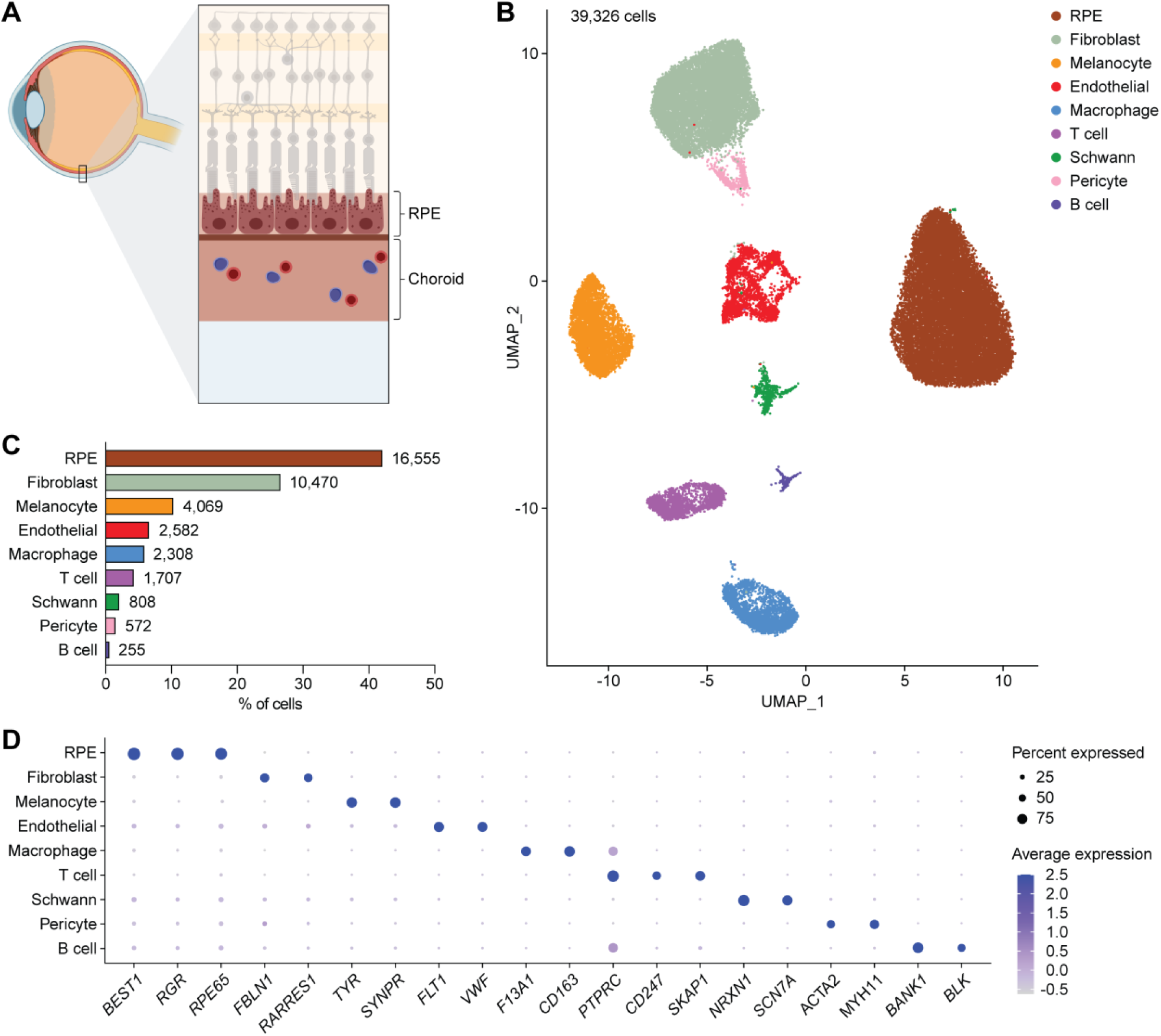
Transcriptional profiles from single-cell multiome sequencing capture major cell types of human RPE and choroid. **(A)** Schematic cross-section of the posterior segment of the eye depicting the anatomic location of the retinal pigment epithelium (RPE) and choroid. **(B)** Uniform manifold approximation and projection (UMAP) plot of the 39,326 human RPE and choroid cells detected by scRNA-seq after quality control filtering, doublet removal, and exclusion of ciliary epithelium and neural retina cells. Tissues from fourteen postmortem eyes from nine donors were profiled. A total of ten RPE and choroid clusters were resolved and assigned to nine cell types. **(C)** Frequency of different RPE and choroid cell types as determined by scRNA-seq. Numbers adjacent to each bar indicate absolute counts. **(D)** Dot plot depicting the normalized RNA expression of selected marker genes by cell type. The color and size of each dot signify the average expression level and percent of expressing cells, respectively.

Using shared barcodes from joint scRNA- and scATAC-seq, we generated chromatin accessibility profiles for seven of the nine cell types annotated by scRNA-seq, excluding pericytes and B cells due to having too few cells for reliable peak calling. The resulting chromatin accessibility profiles clustered by cell type (Figure 2A) and were concordant among donors (Figure S4A), supporting the validity of our multiome. Chromatin accessibility in RPE cells as determined by scATAC-seq was also highly similar to that from published bulk ATAC-seq of human RPE (Figure S3B).^29^ To identify chromatin accessibility peaks for RPE and choroid cell types, we performed MACS2 peak calling on scATAC-seq profiles grouped into pseudo-bulk ATAC replicates. We uncovered a total of 241,538 chromatin accessibility peaks in the RPE and choroid (Figure 2B; Data S2), >50% of which were unique to a single cell type (Figure 2C). These included 16,018 marker peaks significantly enriched in a cell type-specific manner (Figure 2D) and many located near cell type-specific genes (Figure 2E).

**Figure 2.**
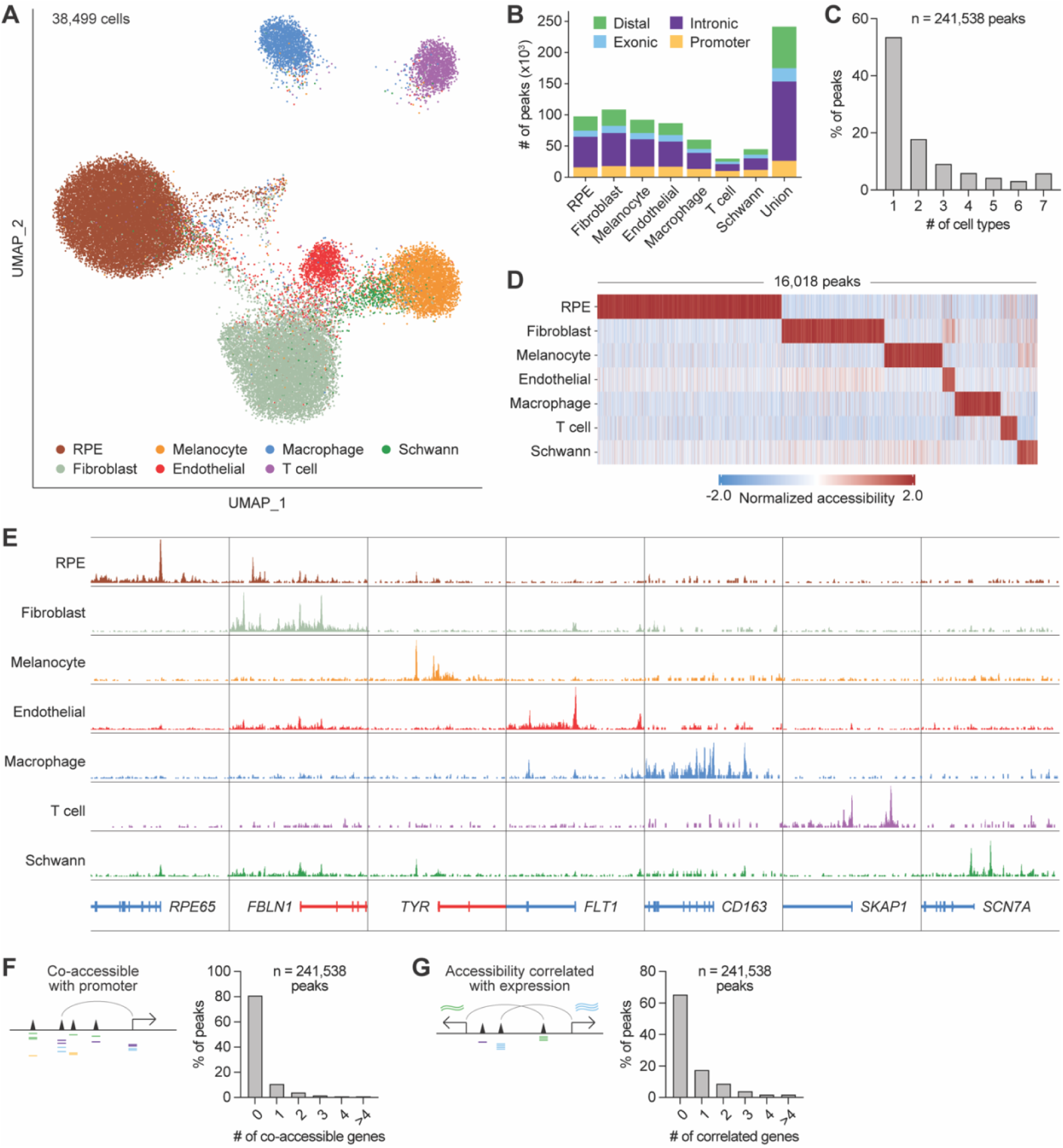
Chromatin accessibility profiles from single-cell multiome sequencing reveal epigenetic landscapes of RPE and choroid cell types. **(A)** UMAP plot of 38,499 human RPE and choroid cells profiled by scATAC-seq. Cell identities were assigned by scRNA-seq. **(B)** Number and characterization of chromatin accessibility peaks in RPE and choroid cell types as determined by scATAC-seq. **(C)** Number of RPE and choroid cell types exhibiting each scATAC-seq peak. **(D)** Heatmap depicting the normalized accessibility of scATAC-seq marker peaks by cell type. Each column represents a marker peak. **(E)** Sequencing tracks of cell type chromatin accessibility near selected marker genes. Each track depicts the aggregate scATAC-seq signal of up to 10,000 cells from the given cell type normalized by the number of reads in transcriptional start site (TSS) regions. Genes transcribed in the sense direction (TSS on the left) are shown in red, while genes in the antisense direction (TSS on the right) are shown in blue. Coordinates for each region: *RPE65* (chr1:68429958-68469959), *FBLN1* (chr22:45472237-45532238), *TYR* (chr11:89147451-89207452), *FLT1* (chr13:28455127-28535128), *CD163* (chr12:7483892-7523893), *SKAP1* (chr17:48410274-48450275), *SCN7A* (chr2:166464246-166524247). **(F)** Number of genes with a co-accessible promoter for each scATAC-seq peak in RPE and choroid cell types. Co-accessible was defined as a correlation score >0.3 between the assayed peak and at least one peak within the promoter. **(G)** Number of genes with correlated expression for each scATAC-seq peak in RPE and choroid cell types. Correlated was defined as a correlation score >0.3 between the accessibility of the assayed peak and the RNA expression of the gene.

Using our multiome, we analyzed subpopulations of RPE and choroid cells. We first examined the two RPE clusters identified by scRNA-seq, which we annotated as macular and peripheral RPE (Figure S2C). Comparing these two clusters revealed 18 differentially expressed genes (Data S3), including higher expression of *SULF1* and *WFDC1* in macular RPE and *KCNIP4* and *VEPH1* in peripheral RPE as previously reported.^23^ We likewise compared chromatin accessibility profiles from macular and peripheral RPE to search for regions exhibiting differential accessibility. While several peaks in our dataset appeared to show greater accessibility in macular or peripheral RPE (Figure S4B), none passed our statistical criteria for differential accessibility (log2 fold change ≥0.5 and false discovery rate [FDR] ≤0.25). We therefore combined macular and peripheral RPE into a single cell type for downstream analyses.

We next conducted preliminary comparisons of RPE and choroid cells from eyes with and without dry AMD. All nine cell types were present in both control and diseased tissues and in roughly the same distribution (Figure S5A). Comparison of control and AMD eyes showed 2-56 differentially expressed genes per cell type, including higher levels of heat shock proteins in RPE, lower levels of the antioxidant enzyme *SOD2* in fibroblasts and vascular endothelial cells, and higher levels of the metabolic regulator *PDK4* in macrophages and pericytes (Figure S5B; Data S4). Macular RPE from AMD eyes also upregulated *ANKRD37*, a gene more responsive to hypoxia than *VEGFA*.^30^ We similarly compared chromatin accessibility between control and AMD cell types but were unable to find any differentially accessible regions in RPE, including when analysis was confined to macular RPE. Across all other cell types, we observed only one differentially accessible peak which was unique to macrophages (Figure S4C).

We further leveraged our multiome to characterize potential gene regulatory interactions in RPE and choroid cell types. Using scATAC-seq, we located candidate enhancers by assessing for peaks whose accessibility correlated with that of a nearby promoter, supporting a regulatory relationship between these elements. This method detected co-accessible promoters for 45,731 (18.9%) of the 241,538 scATAC-seq peaks in our dataset and nominated 63,726 peak-gene pairs as enhancer interactions (Figure 2F; Data S5). We additionally identified candidate enhancers by searching for scATAC-seq peaks whose accessibility correlated with the scRNA-seq expression of a nearby gene, suggesting an effect on gene regulation. Employing this strategy, we found 83,491 RPE and choroid peaks with at least one correlated gene and 161,269 peak-gene pairs (Figure 2G; Data S6).

Finally, we performed motif enrichment analysis using peaks from scATAC-seq to predict transcription factor (TF) activity in RPE and choroid cell types (Figure S6A; Data S7). In agreement with published literature, we saw enrichment of binding motifs for OTX family members in RPE,^31,32^ ATF4 in fibroblasts,^33^ SOX family members in melanocytes,^34^ EBF1 in vascular endothelial cells,^35^ SPI family members in macrophages,^36,37^ ETS family members in T cells,^38^ and NR1H4 in Schwann cells.^39^ Footprinting analysis of scATAC-seq peaks moreover supported TF occupancy and activity by demonstrating motif centers to be protected from Tn5 transposition (Figure S6B).

In summary, our multiome reveals the gene expression and chromatin accessibility landscapes of human RPE and choroid at single-cell resolution. Analyses of this dataset identify marker genes and peaks for RPE and choroid cell types, nominate enhancers and TFs in these tissues, and uncover potential cell type-specific perturbations in AMD.

### H3K27ac HiChIP and Activity-by-Contact map enhancer-gene interactions in the RPE and choroid

The 3D organization of the genome enables enhancers and other regulatory elements to interact with genes and control their expression.^40^ To elucidate 3D enhancer-gene interactions in human RPE and choroid, we performed HiChIP for H3K27ac, a histone mark of active enhancers and promoters (Figure 3A; Figure S7A).^12,41^ H3K27ac HiChIP reads were enriched near peaks from H3K27ac chromatin immunoprecipitation sequencing (ChIP-seq) of human RPE and choroid and an RPE cell line (Figure S7B),^42,43^ supporting the validity of our HiChIP results. This enrichment was not seen near peaks from ChIP-seq for H3K27 trimethylation (H3K27me3), a mark of repressed genes.^44^ H3K27ac HiChIP identified 2,145 loop anchors connected by 1,116 loops, 97.6% of which overlapped a scATAC-seq peak in at least one anchor (Figure 3B; Data S8). Some of these loops linked scATAC-seq peaks to cell type-specific genes (Figure 3C), consistent with the detection of cell type-specific enhancer-gene interactions.

**Figure 3.**
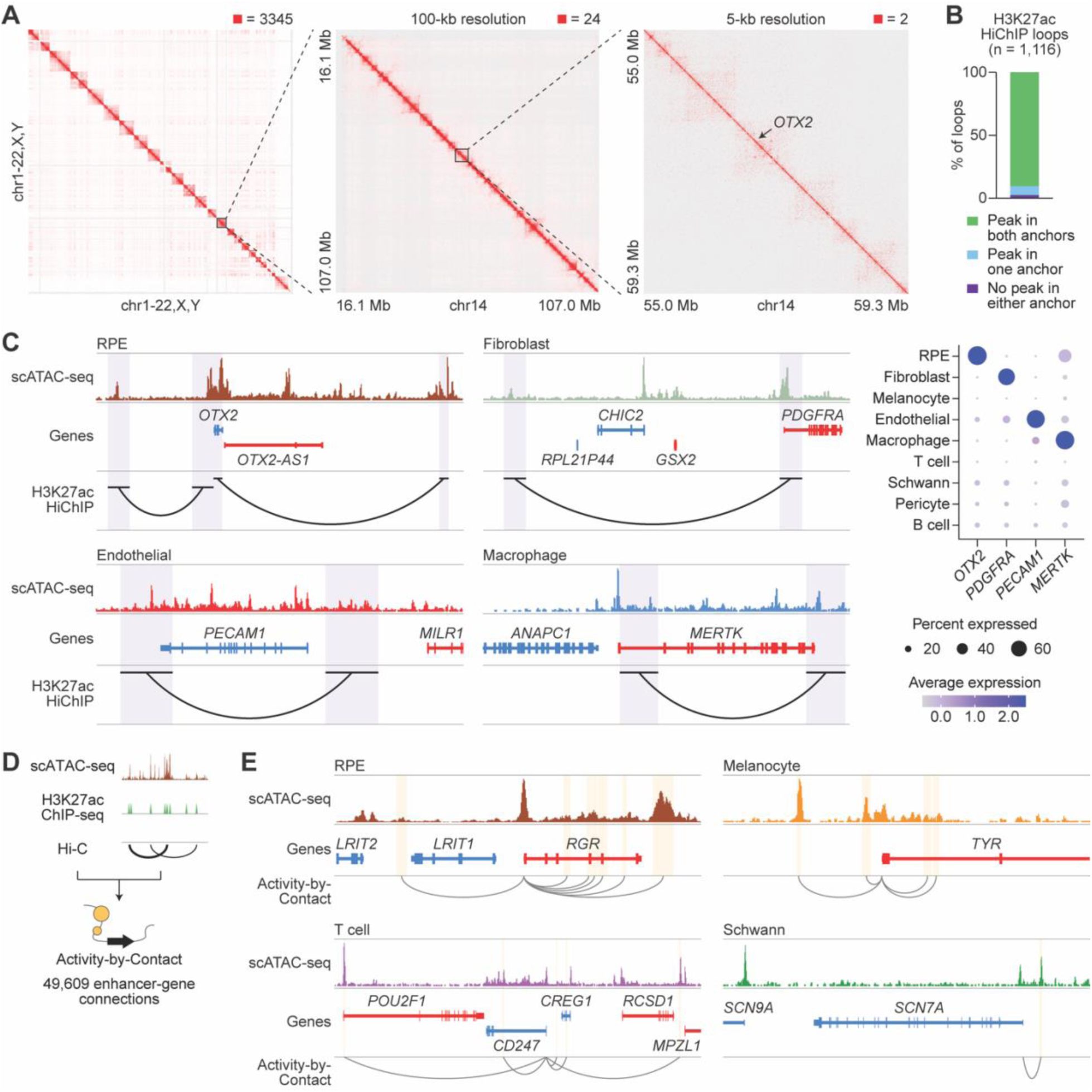
H3K27ac HiChIP and Activity-by-Contact of RPE and choroid identify candidate enhancer-gene interactions. **(A)** 3D interaction maps from human RPE and choroid at whole genome, 100-kb, and 5-kb resolution. Signal was normalized to the square root of coverage. Numbers above each interaction map indicate the maximum signal displayed. **(B)** Overlap of H3K27ac HiChIP loop anchors with scATAC-seq peaks. Overlap was defined as any overlapping bases. **(C)** Left: sequencing tracks of cell type chromatin accessibility and H3K27ac HiChIP loops overlapping selected cell type-specific genes. Genes transcribed in the sense and antisense directions are shown in red and blue, respectively. Regions encompassed by loop anchors are highlighted in purple. Coordinates for each region: RPE (chr14:56661727-57099176), fibroblast (chr4:53876209-54307966), endothelial (chr17:64288587-64467042), macrophage (chr2:111808822-112054161). Right: dot plot depicting the normalized RNA expression of selected cell type-specific genes overlapping H3K27ac HiChIP loops. **(D)** Schematic of the Activity-by-Contact (ABC) model to predict enhancer-gene connections. scATAC-seq reads were combined with public H3K27ac ChIP-seq^42^ and Hi-C^45^ data from human RPE and choroid to calculate ABC scores. **(E)** Sequencing tracks of cell type chromatin accessibility and enhancer-gene connections for selected cell type-specific genes. Enhancers predicted by ABC are highlighted in yellow. Gray arcs depict enhancer-gene connections. Coordinates for each region: RPE (chr10:84222538-84266351), melanocyte (chr11:89159418-89201409), T cell (chr1:167207660-167746132), schwann (chr2:166367444-166513713).

As a complementary approach to find enhancers, we also used the ABC model, which has been shown to accurately identify enhancers and their target genes.^21^ This model weighs activity at candidate sequences by how often they contact gene promoters to predict enhancer-gene connections. To calculate ABC scores, we combined our scATAC-seq reads with public H3K27ac ChIP-seq and Hi-C data from human RPE and choroid.^42,45^ ABC analysis revealed 21,294 enhancers and 49,609 enhancer-gene connections, including many connecting scATAC-seq peaks to cell type-specific genes (Figures 3D and 3E; Data S9).

We compared candidate RPE and choroid enhancers detected by various mapping strategies including 1) co-accessibility with a promoter, 2) correlation with gene expression, 3) H3K27ac HiChIP, and 4) ABC (Figure 4A). These methods preferentially captured different numbers and sizes of enhancer-gene interactions (Figures 4B and 4C), limiting overlap among strategies to 20-47% (Figure S7C). Nonetheless, enhancer targets from all four approaches were enriched for RPE and choroid marker genes (Figure 4D), supporting the authenticity of nominated enhancer-gene pairs. In total, our datasets offer a genome-wide “connectome” of enhancers in human RPE and choroid, detailing the regulatory interactions that govern expression in these tissues.

**Figure 4.**
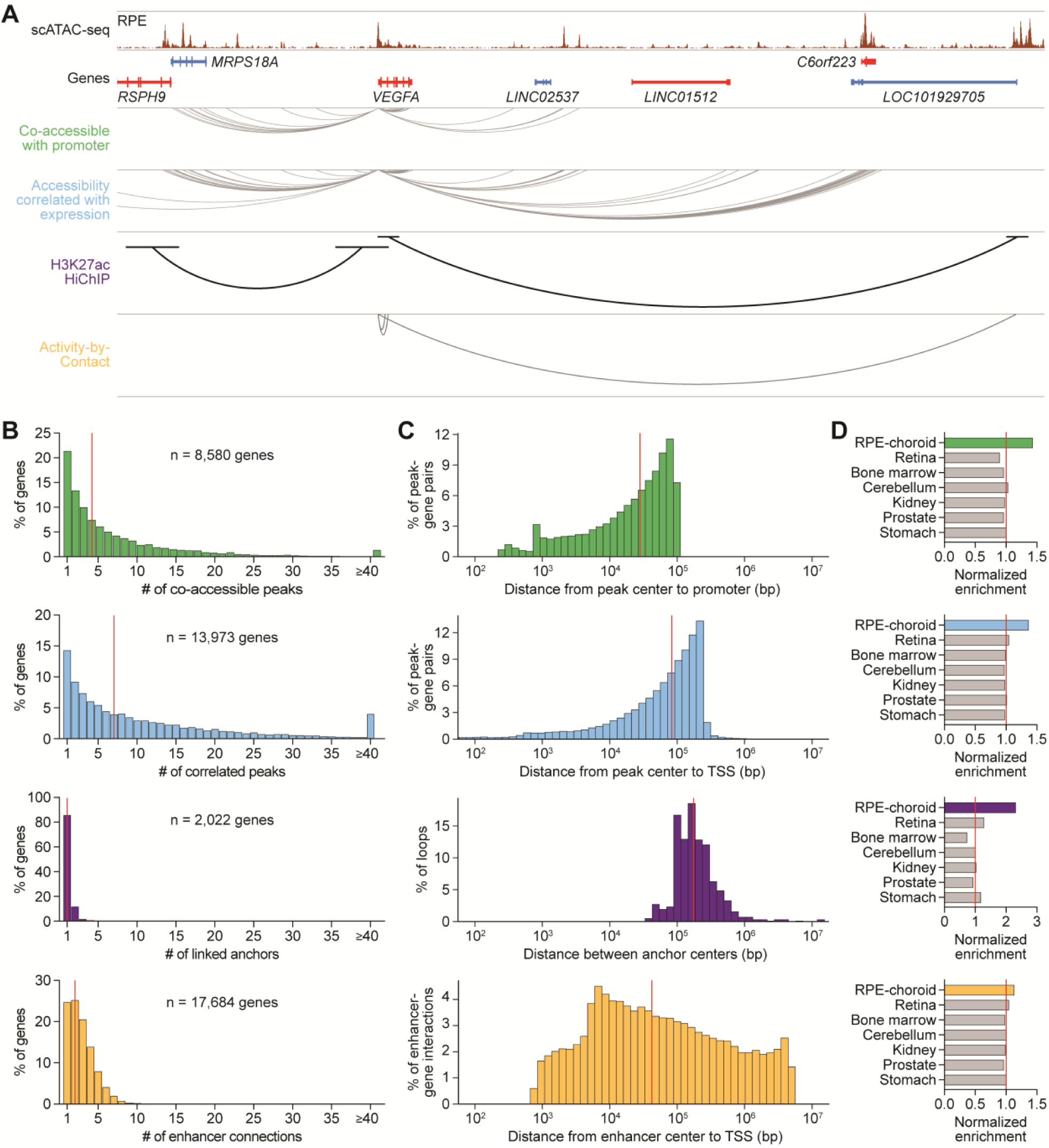
Orthogonal mapping strategies construct a genome-wide connectome of RPE and choroid enhancers. **(A)** Sequencing track of RPE chromatin accessibility and enhancer mapping strategies at the *VEGFA* locus. Candidate sequences were identified as enhancers if 1) they were co-accessible with a promoter, 2) their accessibility correlated with gene expression, 3) they were linked to a gene by H3K27ac HiChIP, or 4) they were connected to a gene by ABC. Genes transcribed in the sense and antisense directions are shown in red and blue, respectively. Gray and black arcs depict locations of *VEGFA* enhancers detected by each strategy. Coordinates for region: chr6:43645888-44088197. **(B)** Number of enhancers per gene for each enhancer mapping strategy. Colors correspond to the four strategies listed in (A). Red lines depict medians. **(C)** Distance between enhancers and their target genes for each enhancer mapping strategy. Red lines depict medians. Analysis of H3K27ac HiChIP was confined to loops containing at least one gene. TSS, transcriptional start site. **(D)** Enrichment of scRNA-seq marker genes among target genes from each enhancer mapping strategy. Enrichment was calculated as the fraction of marker genes represented among enhancer target genes. Marker genes from retina, bone marrow, cerebellum, kidney, prostate, and stomach were obtained from public datasets.^11,68^ Values were normalized to the average in non-ocular tissues (red lines).

### Single-cell multiomics pinpoint cell type targets of variants associated with AMD

Using our RPE and choroid datasets, we sought to interrogate genomic risk loci identified by GWAS of AMD. Previously, a large GWAS by Fritsche et al. found 52 independent AMD-associated variants distributed across 34 loci.^8^ Forty-seven (90.4%) of these variants constitute noncoding or synonymous changes, making them challenging to interpret with transcriptomic data alone. To evaluate AMD risk loci, we first performed linkage disequilibrium expansion on the 52 index signals to capture nearby variants with high frequency of coinheritance. Linkage disequilibrium expansion resulted in a total of 1,998 AMD risk variants (Figure 5A; Data S10), of which 1,964 (98.3%) were noncoding or encoded a synonymous substitution.

**Figure 5.**
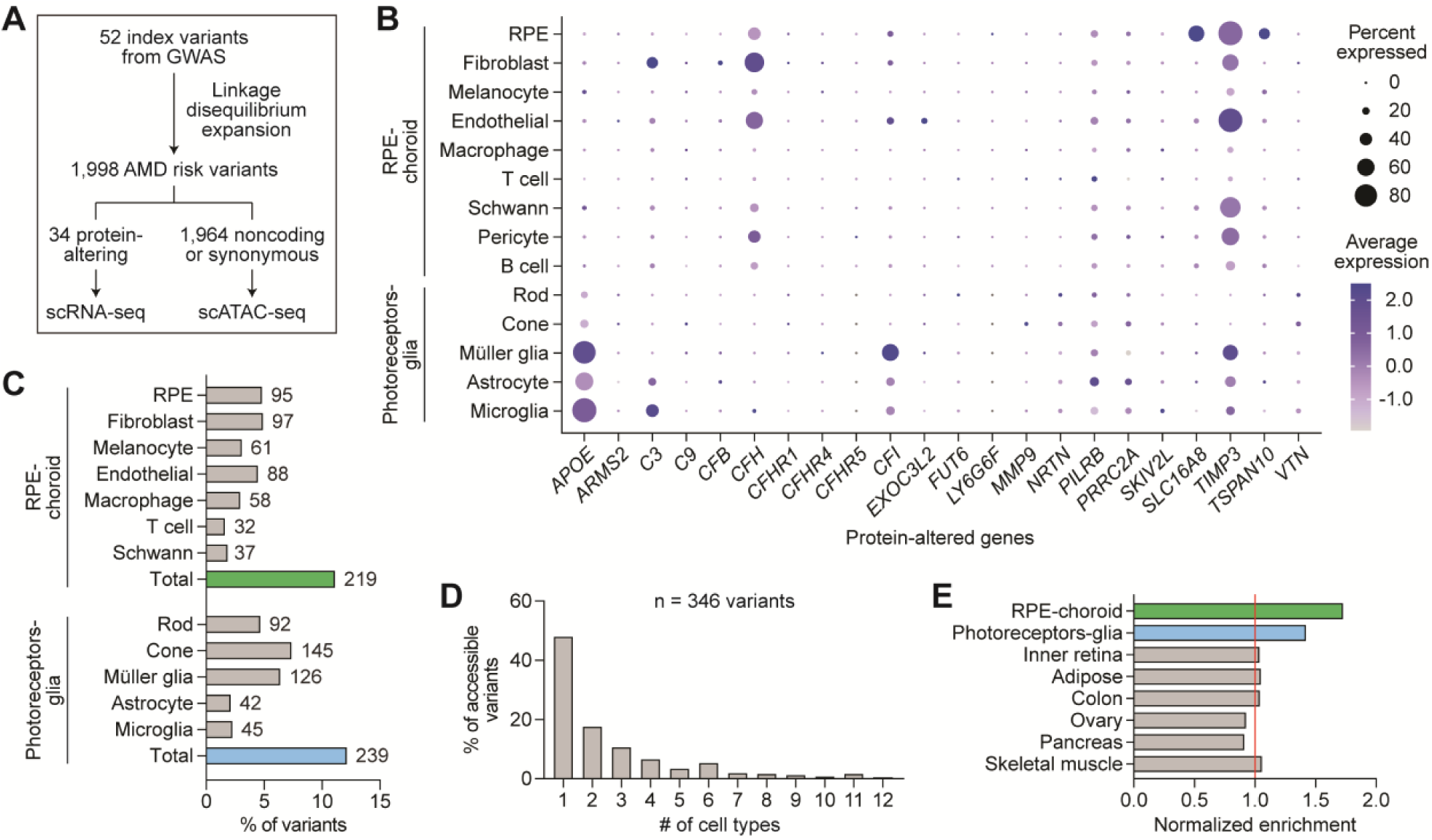
Integration of single-cell multiomics and GWAS uncovers cell type targets of variants associated with AMD. **(A)** Overview of variant selection. Index variants from Fritsche et al.^8^ were subjected to linkage disequilibrium expansion to obtain risk variants for AMD. Coding variants were analyzed using scRNA-seq profiles from RPE, choroid, photoreceptor, and glia cell types to characterize variant expression. Noncoding or synonymous variants were analyzed using scATAC-seq peaks to identify accessible variants. **(B)** Dot plot depicting the normalized RNA expression of genes affected by protein-altering variants in RPE, choroid, photoreceptor, and glia cell types. **(C)** Percent of AMD risk variants accessible in RPE, choroid, photoreceptor, and glia cell types. Numbers adjacent to each bar indicate absolute counts. **(D)** Number of accessible cell types per variant. Only variants accessible in at least one RPE, choroid, photoreceptor, or glia cell type were analyzed. **(E)** Enrichment of noncoding or synonymous AMD risk variants in scATAC-seq peaks from selected tissues. Enrichment was calculated by dividing the number of accessible variants by the number of unique bases in scATAC-seq peaks. Peaks from inner retina, adipose, colon, ovary, pancreas, and skeletal muscle were obtained from public datasets.^11,69^ Values were normalized to the average in non-ocular tissues (red line).

Thirty-four AMD risk variants encoded changes to protein sequences. Using scRNA-seq, we assessed the expression of these protein-altering variants in RPE and choroid cell types, as well as photoreceptor and glia cell types from human retina given their suspected involvement in AMD (Figure 5B).^11^ We also analyzed the expression of four genes (*CFH*, *CFI*, *TIMP3*, *SLC16A8*) suggested by Fritsche et al. to be causal based on rare coding or splice-altering variants.^8^ scRNA-seq demonstrated RPE-specific expression of *SLC16A8* and *TSPAN10* and glia-specific expression of *APOE*, in line with prior reports.^46,47^ Complement genes including *C3*, *CFH*, and *CFI* were most commonly expressed in fibroblasts and vascular endothelial cells, while several genes such as *PILRB* and *TIMP3* were more broadly expressed across cell types.

We additionally used AlphaMissense, a deep learning model that predicts the pathogenicity of single amino acid substitutions,^48^ to score risk variants encoding missense mutations. Of note, 23 (88.5%) of the 26 AMD risk variants in the AlphaMissense database were designated as likely benign (Data S10), suggesting that other variants at these loci might instead drive their association with AMD.

We then examined the 1,964 noncoding or synonymous variants linked to AMD. To identify cell types in which these variants might possess regulatory activity, we intersected their coordinates with scATAC-seq peaks from our dataset as well as from photoreceptors and glia.^11^ This approach uncovered 346 accessible AMD risk variants including 219 in the RPE and choroid (Figure 5C), with most variants accessible in only one or two cell types (Figure 5D). AMD risk variants were enriched in peaks from the RPE, choroid, photoreceptors, and glia (Figure 5E), supporting these cells and tissues as key sites of AMD pathogenesis. Collectively, these analyses elucidate the cell type targets of both coding and noncoding variants implicated in AMD.

### STARR-seq identifies AMD risk variants with enhancer activity in RPE cells

We next performed functional testing of AMD risk variants using STARR-seq, a high-throughput method to screen enhancers.^15,49^ This technique places candidate sequences downstream of a minimal promoter such that active enhancers transcribe themselves, enabling quantification of their regulatory activity. We focused on single-nucleotide polymorphisms (SNPs), representing 89.8% of AMD risk variants, and conducted allele-specific STARR-seq in ARPE-19 cells, a human RPE cell line (Figure 6A).^50^ For each SNP, we synthesized the reference or alternative allele centered in 152 base pairs (bp) of genomic context. The resulting oligonucleotide library was transfected into RPE cells and transcription of each allele compared after 24 hours. Given strong genetic associations between complement genes and AMD, we reasoned that complement activation might modulate the regulatory activity of some SNPs. We thus also conducted STARR-seq in RPE cells supplemented with serum containing activated complement factors to assess SNPs in the context of complement activation.

**Figure 6.**
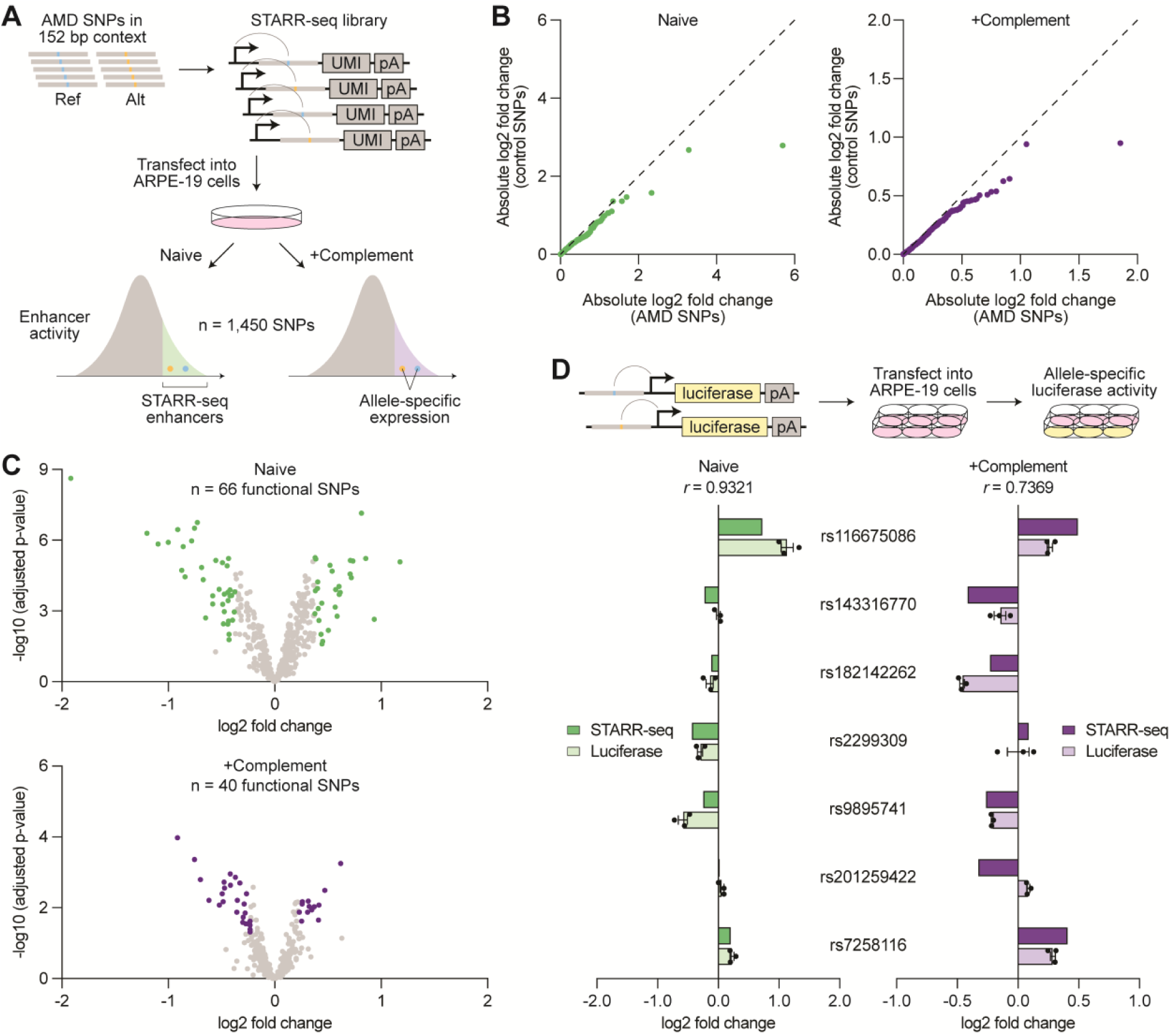
Allele-specific STARR-seq quantifies the regulatory activity of AMD risk variants in RPE cells. **(A)** Schematic of allele-specific STARR-seq. Reference and alternative alleles of single-nucleotide polymorphisms (SNPs) at AMD risk loci were centered in 152 bp of genomic context and placed downstream of a minimal promoter, enabling sequences with enhancer activity to transcribe themselves. The resulting STARR-seq library was transfected into ARPE-19 cells cultured with or without activated complement factors, then transcription of each allele measured. Reference and alternative allele expression were compared with the input library to identify STARR-seq enhancers and with each other to quantify allele-specific activity. **(B)** Quantile-quantile plots comparing the allele-specific expression (alternative/reference) of SNPs at AMD risk loci versus control SNPs randomly selected from the *MAPT* locus. Each dot (n = 357) represents the same quantile of absolute log2 fold change in both groups. Dashed lines depict identical distributions. **(C)** Volcano plots depicting the allele-specific expression (alternative/reference) of AMD-associated SNPs in STARR-seq enhancers. Each dot represents a SNP (n = 462 for Naive, n = 342 for +Complement). Functional SNPs in naive and complement-activated RPE cells are labeled in green and purple, respectively. **(D)** Allele-specific enhancer activity (alternative/reference) of selected SNPs measured by STARR-seq versus a luciferase reporter. Data shown are mean ± SEM. *r* values indicate Pearson correlation coefficients.

STARR-seq demonstrated high reproducibility, with correlation scores of 0.972-0.994 across replicates of the same condition (Figure S8A). An average of 3,890 unique barcodes were detected per oligonucleotide in the input library, while output libraries from naive and complement-activated RPE cells averaged 316 and 272 unique barcodes per oligonucleotide, respectively (Figure S8B). Out of 1,795 total SNPs at AMD risk loci, 1,450 (80.8%) passed quality control filters (≥5 unique barcodes per oligonucleotide in all replicates) and were included in the final analysis (Data S11).

We first located SNPs residing in genomic regions with enhancer activity, which we termed STARR-seq enhancers, by identifying oligonucleotides overrepresented in output libraries relative to the input library (Figure S9A). These regions were significantly enriched for scATAC-seq peaks from RPE and choroid cell types (Figure S9B), consistent with their role as regulatory elements. We uncovered 462 and 342 AMD-associated SNPs within STARR-seq enhancers in naive and complement-activated RPE cells, respectively, and 150 within enhancers active in both conditions (Data S11). We next compared the transcription of reference and alternative alleles to determine SNPs with allele-specific activity. As a control, we performed the same analysis using >350 randomly selected SNPs from the *MAPT* locus implicated in progressive supranuclear palsy,^16^ a condition not expected to involve the RPE. In both naive and complement-activated experiments, AMD-associated SNPs were more likely to impact regulatory activity (Figure 6B), suggesting that some of them were functionally relevant in RPE cells.

We designated SNPs as functional if they 1) were located in a STARR-seq enhancer, and 2) demonstrated a statistically significant (adjusted p-value <0.05) difference in expression between alleles exceeding the 80th percentile of absolute log2 fold change among control SNPs. Using this definition, we found 66 and 40 functional SNPs in naive and complement-activated RPE cells, respectively (Figure 6C). We further tested seven SNPs in a luciferase reporter system and saw strong correlation with STARR-seq (Figure 6D), supporting the validity of our STARR-seq measurements. Altogether, our STARR-seq experiments provide a detailed map of SNP regulatory activity in RPE cells, shedding light on the functional importance of variants linked to AMD.

### Integration of RPE and choroid datasets predicts causal variants in AMD

We conducted two additional analyses to identify AMD risk variants with allele-specific effects on gene expression or chromatin accessibility. Referencing published expression quantitative trait loci (eQTLs) from human RPE and choroid,^46^ we located risk variants whose genotypes were associated with gene expression changes at a FDR ≤0.05 and termed these variants “significant eQTLs.” We also trained deep learning models derived from the ChromBPNet architecture^51–53^ on scATAC-seq profiles from RPE and choroid cell types (Figures S10A and S10B). Using these models, we predicted the effect of reference and alternative alleles for each variant on local chromatin accessibility (Figure S10C) and annotated variants as “high effect” if they exhibited an absolute log2 fold change >0.25 in predicted accessibility at a FDR <0.01 in any cell type.

We then integrated all our methods and datasets to prioritize AMD risk variants, focusing on the 219 variants accessible in RPE and choroid cell types (Figure 7A). Of the 219 variants, 82 (37.4%) were co-accessible with a promoter, 89 (40.6%) showed accessibility that correlated with the expression of a gene, 10 (4.6%) were linked to a gene by H3K27ac HiChIP, 31 (14.2%) were connected to a gene by ABC, 96 (43.8%) were located in a STARR-seq enhancer, 15 (6.8%) were a functional SNP by STARR-seq, 47 (21.5%) were a significant eQTL, and 4 (1.8%) were high effect by ChromBPNet (Figure 7B; Data S10). We designated the 59 variants (26.9%) that met at least three of the eight prioritization criteria as prioritized variants, representing 3.0% of the initial 1,964 noncoding or synonymous variants at AMD risk loci. These variants possessed the strongest evidence for gene regulatory activity in the RPE and choroid, supporting their potential involvement in AMD.

**Figure 7.**
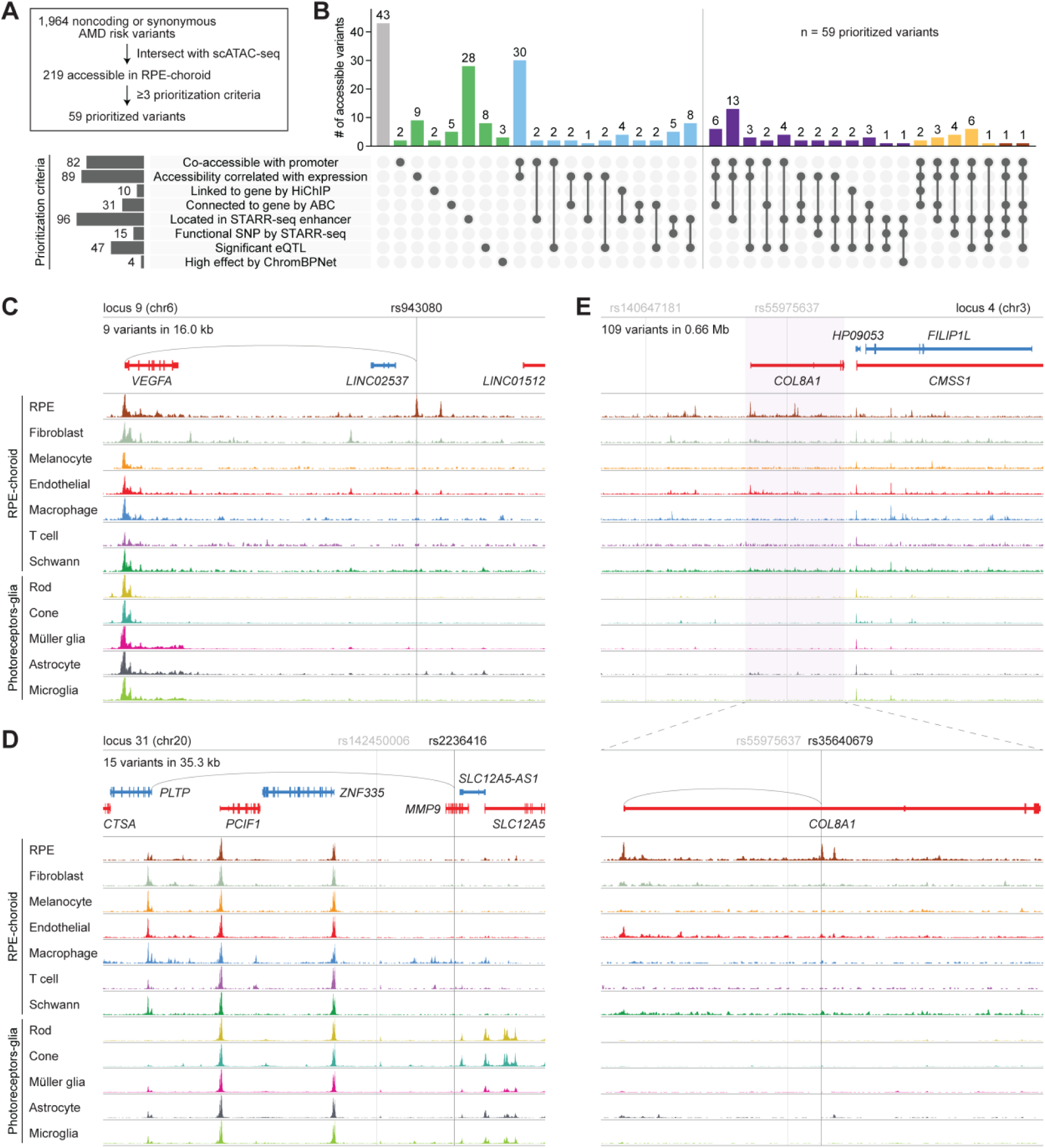
Integrated analysis of RPE and choroid datasets nominates causal variants at AMD risk loci. **(A)** Overview of variant prioritization. Variants accessible in RPE and choroid cell types were designated as prioritized variants if they met at least three of the eight prioritization criteria. **(B)** Upset plot depicting the percent of accessible variants fulfilling each combination of prioritization criteria. Numbers above each bar indicate absolute counts. Prioritized variants are right of the vertical line. **(C-E)** Sequencing tracks of cell type chromatin accessibility at AMD risk loci 9 (C), 31 (D) and 4 (E). Genes transcribed in the sense and antisense directions are shown in red and blue, respectively. Vertical gray lines depict the locations of index variants, and vertical black lines depict the locations of putative causal variants. Gray arcs indicate predicted target genes for putative causal variants. Coordinates for each region: C (chr6:43763764-43897935), D (chr20:45896353-46041679), E top (chr3:99386416-100133954), E bottom (chr3:99630261-99797894).

Finally, we sought to discover causal variants in AMD by systematically evaluating each of the 34 loci from Fritsche et al.^8^ We began by examining the 52 index variants at these loci, 11 (21.2%) of which were accessible in an RPE, choroid, photoreceptor, or glia cell type. Among them was rs943080, the index variant at locus 9, which represents a C-to-T transition on chromosome 6 (Figure 7C). rs943080 was one of two accessible variants and the only prioritized variant at locus 9 and exhibited high accessibility specifically in RPE cells. Although situated between two long intergenic noncoding RNAs, this variant was uniquely co-accessible with the more distal *VEGFA* promoter, and its accessibility correlated with *VEGFA* expression. It also resided in a STARR-seq enhancer, reinforcing its regulatory potential in RPE cells. Our findings predict rs943080 to play a pathogenic role in the RPE by affecting expression of *VEGFA*, the primary therapeutic target for wet AMD.^54^ Agreeing with this, prior bulk RNA-seq of human induced pluripotent stem cell-derived RPE showed significantly lower *VEGFA* levels in cells from a donor homozygous for the rs943080 risk allele than from heterozygous controls.^55^

Nonetheless, most index variants were not accessible in AMD-relevant cell types, suggesting that other variants in linkage disequilibrium might instead be causal. For example, at locus 31 on chromosome 20, the index variant rs142450006 was broadly inaccessible (Figure 7D), arguing against its involvement in this disease. We therefore inspected the 14 variants in linkage disequilibrium with rs142450006 including rs52794457, which encodes a missense mutation in *MMP9*, and rs2236416, a prioritized variant representing an A-to-G transition in an *MMP9* intron. Despite rs52794457 being a protein-altering variant, we were doubtful of its role in AMD given minimal *MMP9* expression in the retina, RPE, and choroid (Figure 4B) and the variant’s designation by AlphaMissense as likely benign.^48^ In contrast, rs2236416 demonstrated accessibility in macrophages, was located in a STARR-seq enhancer, and was uniquely co-accessible with the promoter of phospholipid transfer protein (*PLTP*), a key regulator of lipid and lipoprotein metabolism.^56^ The accessibility of rs2236416 moreover correlated with *PLTP* expression in macrophages, supporting *PLTP* as the target gene for this variant. From these data, we hypothesize that rs2236416 modulates *PLTP* activity in choroidal macrophages, potentially alleviating some of the dysfunctional lipid homeostasis seen in AMD.^57^

We utilized the same strategies to interrogate larger loci such as locus 4 on chromosome 3 (Figure 7E). Fritsche et al. reported two index variants, rs140647181 and rs55975637, at this locus.^8^ However, rs140647181 was not accessible in any AMD-relevant cell types, and rs55975637 failed to meet any criteria for prioritization. Searching the >100 other variants in linkage disequilibrium, we identified rs35640679, which represents a G-to-A transition in an intron of the collagen gene *COL8A1*. rs35640679 was one of only three prioritized variants at locus 4 and the one most accessible in the RPE. This variant was uniquely co-accessible with the *COL8A1* promoter, its accessibility correlated with *COL8A1* expression in RPE cells, and it was predicted by ABC to connect to the *COL8A1* gene. Our results point to rs35640679 as a causal variant at locus 4 and postulate its effect on *COL8A1* in the RPE to confer risk for AMD. More broadly, these examples highlight the utility of our datasets to dissect complex loci implicated in AMD and nominate new targets and mechanisms for this disease.

## DISCUSSION

In this study, we applied a multiomic framework to human RPE and choroid to identify the genes, accessible regions, and regulatory interactions that define AMD-relevant cell types. Using a combination of scRNA-seq, scATAC-seq, H3K27ac HiChIP, ABC, STARR-seq, eQTLs, and ChromBPNet, we then evaluated nearly 2,000 genomic variants associated with AMD and predicted those with the highest pathogenic potential. Our work builds upon prior single-cell transcriptomic^22,26,46^ and bulk epigenomic^29,42,45^ characterizations of the RPE and choroid by providing a comprehensive and publicly accessible (https://eyemultiome.su.domains/) resource to study cell types in these tissues. Our datasets may moreover offer insights into diseases of the RPE and choroid beyond AMD, such as central serous chorioretinopathy, polypoidal choroidal vasculopathy, and choroidal melanoma.

Research on AMD has been hindered by a lack of animal and cell culture models that accurately recapitulate the condition’s complex pathophysiology. We addressed this challenge by directly profiling human ocular tissues from donors with or without AMD, leveraging a single-cell approach to resolve changes in individual cell types. Using scRNA-seq, we performed preliminary comparisons of control and diseased eyes and uncovered several differentially expressed genes implicating putative mechanisms for AMD. For instance, we measured lower *SOD2* levels in fibroblasts and endothelial cells from AMD donors, suggesting reduced capacity of these cells in affected eyes to counter oxidative stress. This finding may inform therapeutic efforts to upregulate antioxidant activity in AMD by proposing target cell types for such treatments.^58,59^ In contrast, we detected only one scATAC-seq peak in the RPE and choroid that was differentially accessible in AMD. While global reduction in RPE accessibility during AMD has been described,^29^ we did not observe this phenomenon in our scATAC-seq data. Rather, our study was more consistent with Orozco et al., who found no significant differences in retina, RPE, or choroid accessibility between control and AMD donors.^24^

We likewise nominated causal genes for AMD by integrating our multiomic analysis of RPE and choroid with results from GWAS. Among the candidates we prioritized was *PLTP*, whose activity we localized to macrophages. Although *PLTP* expression has been assayed in RPE cell lines,^60,61^ its function in choroidal macrophages has not been explored and may represent a therapeutic target for AMD. Previous proteomics experiments showed that *PLTP* is significantly elevated in the plasma of AMD patients,^60^ supporting the possibility that *PLTP* upregulation might facilitate AMD progression. In mice, genetic deficiency of *Pltp* decreased, and overexpression of *Pltp* increased, the occurrence of atherosclerotic plaques,^62,63^ which share some histologic features with drusen such as the presence of lipid-laden macrophages.^64^ Examination of human atherosclerotic plaques additionally demonstrated high *PLTP* levels in macrophages but minimal signal in normal vessels,^65^ further linking macrophage expression of *PLTP* to these pathologic lesions. It is conceivable that increased *PLTP* activity in choroidal macrophages could favor lipid accumulation in adjacent tissues similar to that seen in atherosclerosis, leading to the formation of lipid-rich drusen beneath the RPE. By modulating *PLTP* expression, rs2236416 may combat these processes, protecting against the development of AMD.

### Limitations of the study

Limitations of this work include the low numbers of choroidal pericytes and B cells sampled, precluding identification of scATAC-seq peaks for these cell types. Our study also profiled only four eyes with AMD, which may have reduced our ability to detect differentially expressed genes and accessible regions between AMD and control tissues. While we were able to annotate macular RPE cells using published marker genes,^23^ macular versus peripheral origin for choroid cell types could not be determined. Future expansion of our atlas with additional cells and donors would help address these shortcomings and may uncover more subtle gene expression and chromatin accessibility findings than presented here.

We further noted that methods to predict variant-target gene interactions were often not in agreement, highlighting the complexity of interpreting noncoding variants and value of a multiomic approach. In line with our observations, Fulco et al. reported only modest concordance between H3K27ac HiChIP and ABC modeling in identifying enhancer-gene pairs, with the latter more closely aligned to data from CRISPR interference.^21^ Mostafavi et al. also showed that there is limited overlap between significant variants from GWAS and eQTL studies due to systematic biases in how these variants are discovered.^66^ Using STARR-seq, we sought to test the functional activity of variants, although chromatin contexts in this episome-based assay may have differed from physiological conditions.^67^ Future gene or base editing experiments targeting individual variants in pertinent RPE and choroid cell types will be important to validate our results and fully elucidate how pathogenic variants contribute to AMD.

## ACKNOWLEDGEMENTS

The authors are grateful to Lions VisionGift, Lions Gift of Sight, and the donors who made this study possible. Computing for this project was performed on the Sherlock cluster, a resource provided and maintained by the Stanford Research Computing Center. This work was supported by National Institutes of Health (NIH) grant RM1-HG007735 (to H.Y.C.). S.K.W. was supported by a National Eye Institute training grant (T32EY027816) and the Knights Templar Eye Foundation. H.Y.C. was an Investigator of the Howard Hughes Medical Institute.

## AUTHOR CONTRIBUTIONS

S.K.W., J.L., Y.L., N.W., and H.Y.C. conceived the project and designed experiments. S.K.W., J.L., Y.C., and Y.Z. performed all experiments. S.K.W., J.L., S.N., and R.K. performed data analyses. A.K. Y.L., N.W., and H.Y.C. supervised the work. S.K.W., J.L., Y.L., N.W., and H.Y.C. wrote the manuscript with input from all authors.

## STAR METHODS RESOURCE AVAILABILITY

### Lead contact

Further information and requests for resources and reagents should be directed to and will be fulfilled by the lead contact, Dr. Howard Y. Chang.

### Materials availability

Due to the limiting nature of primary samples, human tissues used in this study are not available upon request. All other unique reagents generated in this study are available from the lead contact with a completed materials transfer agreement.

### Data and code availability

Raw and processed scRNA-seq, scATAC-seq, HiChIP, and STARR-seq data from this study have been uploaded to Gene Expression Omnibus (GEO) under the accession numbers GSE262151 and GSE289703. Code used for scRNA- and scATAC-seq analyses is available at https://github.com/seankwang/RPEchoroid-multiome. ChromBPNet models are available at https://doi.org/10.5281/zenodo.14031498. A web page summarizing our data is available at https://eyemultiome.su.domains/.

## EXPERIMENTAL MODEL AND SUBJECT DETAILS

Postmortem adult human eyes were procured from consented donors by Lions VisionGift (Portland, OR, USA) or Lions Gift of Sight (St. Paul, MN, USA) under protocols approved by the Eye Bank Association of America. De-identified ophthalmic records for each donor eye were reviewed by an ophthalmologist (S.K.W.), and a diagnosis of dry (non-exudative) AMD was accepted if 1) drusen or geographic atrophy were reported in the macula exam, 2) the donor was taking AREDS2 supplements,^70^ and 3) there was no history to indicate conversion to wet AMD, such as blood in the macula or treatment with anti-angiogenic agents. Dry AMD eyes were deemed advanced stage if geographic atrophy was present and early or intermediate stage if not. Only eyes without ocular disease other than cataract or dry AMD were used. Whole RPE-choroid complexes were dissected out from the adjacent neural retina and sclera and flash-frozen in liquid nitrogen with a maximum death-to-preservation interval of 16 hours. Frozen tissues were shipped to Stanford University for processing. Donor information is listed in Table S1.

## METHOD DETAILS

### Nuclei isolation

Nuclei were isolated from frozen RPE and choroid using the Omni-ATAC protocol (https://doi.org/10.17504/protocols.io.6t8herw) with several key modifications.^25^ For each sample, approximately one-third of the frozen RPE-choroid complex was cut into pieces and Dounce homogenized in cold homogenization buffer containing 0.3% IGEPAL CA-630 in the presence of protease and RNase inhibitors. Released nuclei were passed through a 40-micron filter, purified via two rounds of iodixanol gradient centrifugation, and washed with ATAC resuspension buffer containing RNase inhibitor, 0.1% Tween-20, and 2% bovine serum albumin. During collection of nuclei from iodixanol gradients, care was taken to minimize transfer of pigmented debris. Purified nuclei were then passed through a second 40-micron filter, treated with digitonin according to the 10x Genomics demonstrated protocol for complex tissues (CG000375, Rev. B), passed through a 10-micron filter to remove aggregates, and resuspended in diluted nuclei buffer. Nuclei were counted using a manual hemocytometer to achieve a targeted recovery of 5,000 nuclei per sample. A detailed protocol is provided in Protocol S1.

### scRNA- and scATAC-seq library generation

Joint scRNA- and scATAC-seq libraries were prepared using the 10x Genomics Single Cell Multiome ATAC + Gene Expression kit according to manufacturer’s instructions. Libraries were sequenced with paired-end 150-bp reads on an Illumina NovaSeq 6000 or NextSeq 550 to a target depth of 250 million read pairs per sample. For two eyes (LGS6OD and LGS8OS), a second set of scRNA- and scATAC-seq libraries were generated to increase the number of nuclei profiled.

### HiChIP library generation

H3K27ac HiChIP was performed as previously described using a total of ∼1.5 million nuclei from eight frozen eyes (LGS1 OS, LGS2 OS, LGS3 OS, LGS4 OD, LGS4 OS, LGS5 OS, LVG1 OD, and LVG1 OS).^11^ Following isolation, nuclei were pooled, washed with nuclei isolation buffer from the diploid chromatin conformation capture (Dip-C) protocol,^71^ and fixed with 2% paraformaldehyde at room temperature for 10 minutes. Fixed nuclei were washed twice with cold 1% bovine serum albumin in phosphate-buffered saline before resuspension in 0.5% sodium dodecyl sulfate and resumption of the published HiChIP protocol.^12^ Digestion was performed using the MboI restriction enzyme. Sonication was conducted using a Covaris E220 with five duty cycles, peak incident power of 140, and 200 cycles per burst for four minutes. Chromatin immunoprecipitation (ChIP) was achieved using a validated anti-H3K27ac antibody. The HiChIP library was sequenced with paired-end 75-bp reads on an Illumina NextSeq 550 to a target depth of 500 million read pairs.

### STARR-seq plasmid library generation

The input plasmid library for STARR-seq was generated as previously described with minor modifications.^15,49^ Briefly, oligonucleotide sequences containing the reference or alternative allele for each SNP centered in 152 bp of genomic context and flanked by 5’ and 3’ PCR adaptors were ordered from Agilent. This was performed for all SNPs at AMD risk loci, as well as 500 control SNPs randomly selected from the *MAPT* locus.^16^ Using the KAPA HiFi HotStart ReadyMix system, 2.5 ng of pooled oligonucleotides were PCR amplified in a 50 µL reaction for 13 cycles with primers STARR-seq-1F and −1R (Table S3) to add a 10-nucleotide degenerate barcode and adapters for cloning. PCR-amplified oligonucleotides were cloned into the hSTARR-seq_ORI vector backbone (Addgene #99296)^72^ between the origin-of-replication (ORI) and polyadenylation sequences using Gibson assembly, followed by transformation into MegaX DH10B T1R electrocompetent cells (Thermo Fisher Scientific). A total of ten Gibson assemblies and transformations were carried out, and all transformed *E*. *coli* cells were pooled and grown in four liters of LB media until harvest at an optical density of 1.0. The purified input plasmid library was drop dialyzed before use.

### ARPE-19 cell culture

ARPE-19 cells (CL-0026, Procell) were maintained at 37°C and 5% CO_2_ in DMEM medium (Thermo Fisher) supplemented with 10% fetal bovine serum (Gibco) and 1% penicillin-streptomycin (Gibco). For complement stimulation, cells were additionally cultured for 18 hours until collection with 10% complement competent human serum (S1-LITER, EMD Millipore), a source of activated complement factors.^73^ In each experiment, a total of 10 million ARPE-19 cells were transfected with 10 μg of the STARR-seq plasmid library using a Lonza Nucleofector. Four biological replicates for each condition were performed, and cells were harvested after 24 hours.

### STARR-seq RNA library preparation

Harvested ARPE-19 cells were subjected to total RNA isolation using an M5 HiPer Universal RNA Mini Kit (Mei5bio) followed by mRNA isolation using oligo(dT) Dynabeads (Invitrogen). Isolated mRNA was subsequently treated with TURBO DNase (Thermo Fisher Scientific) and purified using RNAClean XP beads (Beckman Coulter). Target-specific first strand cDNA synthesis was performed using SuperScript III reverse transcriptase (Invitrogen) with 1.5 µg of mRNA per reaction and primer RT (Table S3) to reverse transcribe plasmid mRNA from all cells. The resulting cDNA was treated with RNases A and H, pooled, and underwent 16 cycles of PCR using the KAPA HiFi HotStart ReadyMix system with 5 μL of cDNA template per 50 μL reaction and primers STARR-seq-2F and −2R (Table S3) to amplify target sequences. For indexing, 16 additional cycles of PCR were performed using the TruSeq universal adapter and one of four indexing primers (Table S3) with four technical replicates per index. Final STARR-seq output libraries were purified using AMPure XP beads (Beckman Coulter) and sequenced with paired-end 150-bp reads on an Illumina HiSeq 4000 to ∼3 Gb per library.

To sequence the STARR-seq input library, 10 ng of plasmid DNA per reaction was PCR amplified for 16 cycles using primers STARR-seq-2F and −2R, followed by 16 additional cycles of PCR using the TruSeq universal adapter and one of four indexing primers. Products from four technical replicates were pooled, purified, and sequenced as above.

### Luciferase reporter assay

Oligonucleotides containing the reference and alternative alleles for tested variants were individually cloned into the pGL4.23[luc2/minP] vector (Promega) upstream of the minP promoter and firefly luciferase reporter gene. Each reporter plasmid was then co-transfected with a Renilla luciferase control plasmid (Promega) into ARPE-19 cells using Lipofectamine 3000 (Invitrogen). The regulatory activity of reference and alternative alleles was compared using the Dual-Luciferase Reporter 1000 Assay System (Promega). Twenty-four hours after transfection, cells were washed with PBS and lysed with 200 μL of 1x Passive Lysis Buffer for 10 minutes. To assay firefly luciferase activity, 20 μL of lysate was mixed with 100 μL of Luciferase Assay Reagent II in a 1.5 mL centrifuge tube and luminescence measured using a GloMax 20/20 luminometer (Promega). To assay Renilla luciferase activity, 100 μL of Stop & Glo Reagent was added to the reaction and luminescence again measured. Firefly luciferase activity was normalized to Renilla luciferase activity to control for transfection efficiency and cell number. Three biological replicates were performed for each sequence.

## QUANTIFICATION AND STATISTICAL ANALYSIS

### scRNA- and scATAC-seq data preprocessing and quality control

Demultiplexed FASTQ files from scRNA- and scATAC-seq were inputted into the Cell Ranger ARC pipeline (version 2.0.0) from 10x Genomics to create barcoded count matrices of gene expression and chromatin accessibility data. For each sample, count matrices were loaded in ArchR (version 1.0.2) and selected for barcodes present in both scRNA-seq and scATAC-seq datasets.^74^ Samples in ArchR were quality control filtered for nuclei with 200-10,000 RNA transcripts, <1% mitochondrial reads, <5% ribosomal reads, transcriptional start site (TSS) enrichment >4, and >1,000 ATAC fragments. Quality control filtered nuclei then underwent automated double removal using the filterDoublets function.

### scRNA-seq data analysis

scRNA-seq data from nuclei remaining after quality control filtering and automated doublet removal were analyzed using Seurat (version 3.1.5).^75^ For each sample, gene expression counts were normalized, scaled, and subjected to graph-based clustering using the top 30 principal components at a resolution of 0.3. Marker genes for each cluster were identified using the FindAllMarkers function with a log2 fold change ≥1 and fraction ≥0.6, and cluster identities were manually annotated based on their expression of genes from published scRNA-seq studies of the human eye.^11,22,26–28^ Clusters with no detected marker genes or expressing canonical marker genes from multiple cell types (i.e. putative doublets or multiplets) were removed, after which the dataset was re-clustered using the same parameters. Processed samples were subsequently merged into a single Seurat object, batch corrected using Harmony (version 1.0),^76^ and subjected to graph-based clustering using the top 30 principal components at a resolution of 0.3. Marker genes for each cluster in the merged dataset were identified as above, and clusters with no detected marker genes or expressing canonical marker genes from multiple cell types were removed, after which the dataset was re-clustered. For the final dataset, clusters corresponding to pigmented ciliary epithelium, nonpigmented ciliary epithelium, and photoreceptors were removed, and clusters corresponding to macular and peripheral RPE were combined.

Pairwise comparison of cell type subpopulations was performed using the FindMarkers function with a log2 fold change ≥0.5 and fraction ≥0.3. Genes with an adjusted p-value <0.05 were considered differentially expressed. scRNA-seq data for photoreceptor and glia cell types were previously generated and are available under GEO accession number GSE196235.^11^

### scATAC-seq data analysis

scATAC-seq data were assigned to cell type identities from scRNA-seq using shared barcodes and analyzed using ArchR (version 1.0.2).^74^ For each cell type, pseudo-bulk ATAC replicates were generated using the addGroupCoverages function with default parameters, which creates five replicates of up to 500 cells each for peak calling. Pericytes and B cells were excluded due to having too few cells for reliable analysis. Chromatin accessibility peaks on chromosomes 1-22 and X and outside of blacklist regions for each cell type were called using the addReproduciblePeakSet function and MACS2 (version 2.2.9.1).^77,78^ Marker peaks were identified using the getMarkerFeatures function with a log2 fold change ≥0.5 and FDR ≤0.1 as determined by Wilcoxon pairwise comparisons. Differentially accessible peaks between cell type subpopulations were likewise identified using the getMarkerFeatures function with a log2 fold change ≥0.5 and FDR ≤0.25. Promoters were defined as regions 2,000 bp upstream to 100 bp downstream of a TSS, and accessible promoters were defined as promoters containing at least one scATAC-seq peak. Co-accessible peaks were identified using the getCoAccessibility function with a correlation cutoff of 0.3 and resolution of 1. Peaks were considered co-accessible with a promoter if co-accessible with at least one peak within the promoter. Genes correlated with scATAC-seq peaks were identified using the getPeak2GeneLinks function integrating barcode-matched scRNA-seq data with a correlation cutoff of 0.3, resolution of 1, and the default maximum distance of 250 kb. scATAC-seq peaks for photoreceptor and glia cell types were previously reported with data available under GEO accession number GSE196235.^11^

### Sequencing tracks

bigWig files were generated in ArchR using the getGroupBW function and were visualized on the WashU Epigenome Browser.^74,79^ A tile size of 100 bp, maximum cell number of 10,000, and normalization by total number of reads in TSS regions were used. Sequencing tracks from retina cell types were similarly generated using scATAC-seq data from Wang et al.^11^ All tracks were aligned to the hg38 reference genome.

### Motif enrichment analysis

TF motif enrichment analysis was performed on scATAC-seq peaks using the peakAnnoEnrichment function in ArchR with default parameters and position frequency matrices from Cis-BP.^74,80^ Footprinting analysis of TFs was conducted using the getFootprints function. To correct for Tn5 insertion bias, the Tn5 insertion signal was subtracted from footprinting signals before plotting.

### ChIP-seq data processing

ChIP-seq data from human RPE and choroid and RPE-1 hTERT cells were obtained from GEO accession numbers GSE137311 and GSE210402.^42,43^ Sequencing reads were trimmed using Trimmomatic (version 0.39), aligned to the hg38 genome using Bowtie2 (version 2.3.4.1), filtered for a MAPQ score >10 using SAMtools (version 1.16.1), and subjected to PCR duplicate removal using Picard MarkDuplicates.^81–83^ Bam files from replicates were combined into a single bam file using SAMtools merge. Peak calling was performed using MACS2 (version 2.2.9.1) in comparison to input controls with a q-value of 0.05 to generate narrowPeak files.^77^

### HiChIP data analysis

HiChIP sequencing files were processed using the HiC-Pro pipeline (version 2.11.0) to remove duplicate reads, assign reads to MboI restriction fragments, filter for valid interactions, and generate binned interaction matrices.^84^ HiChIP read coverage around ChIP-seq peaks was calculated using the plot_chip_enrichment.py script from Dovetail Genomics with the HiChIP valid pairs bam file and ChIP-seq narrowPeak files as input. Filtered read pairs from HiC-Pro were converted into .hic files and analyzed using HiCCUPS from the Juicer pipeline (version 1.8.9) to call H3K27ac HiChIP loops.^85^ HiChIP interaction maps depicting all valid interactions identified by HiC-Pro were visualized using Juicebox (version 1.11.08).^86^

### Activity-by-Contact

Activity-by-Contact (ABC) was performed using pseudo-bulk scATAC-seq data, H3K27ac ChIP-seq data from human RPE and choroid (GEO accession number GSE137311),^42^ and Hi-C data from human RPE and choroid (GEO accession number GSE235726).^45^ scATAC-seq reads from the Cell Ranger ARC pipeline for all RPE and choroid cell types except pericytes and B cells were aggregated to generate a fragments file that was converted to tagAlign format. The ABC model (version 1.1.2) was then run on the tagAlign file, H3K27ac ChIP-seq bam files, and combined .hic file at a resolution of 5 kb to calculate ABC scores.^21^ Enhancer-gene connections were defined using default parameters as interactions with an ABC score >0.025 for which the enhancer and TSS were at least 500 bp apart.

### Marker gene enrichment in enhancer targets

Enrichment of enhancer targets for scRNA-seq marker genes was calculated by determining the fraction of marker genes represented among enhancer target genes. For each type of tissue, a randomly selected set of 457 marker genes was tested, corresponding to the number of marker genes identified in RPE and choroid (Data S1). scRNA-seq marker genes from retina were obtained from Wang et al.^11^ scRNA-seq marker genes from non-ocular tissues were obtained from Han et al.^68^

### Variant selection and linkage disequilibrium expansion

Linkage disequilibrium expansion was performed on the 52 index variants from Fritsche et al.^8^ to obtain risk variants for AMD. LDlinkR was first used to identify GRCh38 variants from the 1000 Genomes Project phase 3 genotypes of all populations (“ALL”) correlated (R^2^ >0.5) with an index variant.^87,88^ For index variants not in the 1000 Genomes reference panel, we substituted with the variant in that locus or sublocus possessing the next highest posterior probability to be causal as calculated by Fritsche et al. After running LDlinkR, we combined our list with the 95% credible set of variants from Fritsche et al., a set of 1,343 variants predicted to contain the causal variant for each of the 52 index signals with >95% probability. Filtering out duplicates resulted in a final list of 1,998 AMD risk variants.

### AlphaMissense scoring

AlphaMissense was used to predict the pathogenicity of missense variants.^48^ Variants were classified as “likely benign” for scores <0.34, “likely pathogenic” for scores >0.565, and “ambiguous” otherwise.

### Variant enrichment in scATAC-seq datasets

Enrichment of variants in scATAC-seq datasets was calculated by counting the number of variants accessible in scATAC-seq peaks, then dividing by the number of unique bases in scATAC-seq peaks. For each type of tissue, peaks from constituent cell types were merged. Peaks from retina cell types were obtained from Wang et al.^11^ Peaks from non-ocular cell types were obtained from the Cis-element Atlas.^69^ Inner retina was composed of horizontal, bipolar, amacrine, and retinal ganglion cells. Adipose was composed of adipocytes and mesothelial cells. Colon was composed of colonic goblet, colon epithelial, and colon smooth muscle cells. Ovary was composed of luteal and general smooth muscle cells. Pancreas was composed of acinar, alpha, beta, delta/gamma, and ductal cells. Skeletal muscle was composed of skeletal myocytes, satellite cells, and skeletal muscle fibroblasts.

### STARR-seq data analysis

FASTQ files from STARR-seq were merged into single amplicons using FLASH (version 1.2.11)^89^ and mapped to oligonucleotide sequences using BWA-MEM (version 0.7.12).^90^ Only primary alignment reads that matched the GCTAGCCATTGCGTGTAGAG adapter sequence and with a barcode length of exactly 10 nucleotides were analyzed. Oligonucleotides associated with <5 unique barcodes in any replicate were discarded. For each sample, the number of unique barcodes detected per oligonucleotide was determined. Unique barcode counts were subsequently normalized and compared across samples using DESeq2.^91^ To identify enhancers, counts for reference and alternative versions of each oligonucleotide were combined. STARR-seq enhancers were then defined as oligonucleotides exhibiting an absolute fold change >1.5 (log2 fold change >0.585) and adjusted p-value <0.05 in output libraries relative to the input library.^92^ To identify functional SNPs, the expression of alternative alleles relative to reference alleles was calculated using the mpralm function from mpra (version 1.20.0).^93^ Functional SNPs were defined as SNPs located in STARR-seq enhancers exhibiting an adjusted p-value <0.05 and absolute log2 fold change exceeding the 80^th^ percentile of absolute log2 fold change among control SNPs.

### eQTL analysis

Human RPE-choroid cis-eQTLs (within 1 Mb of gene) were obtained from the Eye eQTL Browser (http://eye-eqtl.com/).^46^ Each of the 1,998 AMD risk variants and their merged rsIDs from dbSNP (https://www.ncbi.nlm.nih.gov/snp/) were searched in the online database. eQTLs with a FDR ≤0.05 in bulk macular or non-macular RPE-choroid were considered significant.

### ChromBPNet model training and variant scoring

scATAC-seq reads from the Cell Ranger ARC pipeline were aggregated by cell type to generate cell type-specific fragments files. We used ChromBPNet v0.1.3 (https://github.com/kundajelab/chrombpnet) and TensorFlow v2.8.0 in Python 3.8.13 to train ChromBPNet models for each cell type.^51–53^ Input peaks for ChromBPNet were called using the ENCODE ATAC-seq pipeline (https://github.com/ENCODE-DCC/atac-seq-pipeline) v1.10.0 and defined as the overlap.optimal_peak.narrowPeak peak set. Model training was then performed using the chrombpnet pipeline command with the scATAC_dermal_fibroblast_bias.h5 bias model (https://zenodo.org/records/7443683/files/bias_models.zip). Five models were trained for each cell type corresponding to the five disjoint training folds specified at https://zenodo.org/records/7443683/files/folds.zip. For fold 0, we ran the chrombpnet contribs_bw command on the counts output within peak regions for each cell type to obtain contribution score bigWig files and ran the chrombpnet modisco_motifs command to summarize motifs using TF-MoDISco.^94^ Models, counts interpretation bigWig files, and TF-MoDISco outputs are available at https://doi.org/10.5281/zenodo.14031498.

To score variants, we followed methods similar to https://github.com/kundajelab/retina-models/tree/main/notebooks.^11^ Briefly, we centered the input window at the variant and obtained the log2 fold change in predicted counts between the reference and alternative alleles for each cell type-specific ChromBPNet model. We averaged the log2 fold change over the five model folds for each variant and cell type. To obtain p-values, we performed one-sided Poisson tests of the predicted alternative allele count with the rate parameter set to the predicted reference allele count (counts averaged over five folds). We then combined p-values for each variant across cell types with Fisher’s method and performed Benjamini-Hochberg correction. Variants with an absolute fold-averaged log2 fold change >0.25 and FDR <0.01 were annotated as high effect. Base importance tracks were visualized using Logomaker.^95^

## SUPPLEMENTAL INFORMATION

**Figure S1.** Single-cell RNA- and ATAC-seq quality control metrics, Related to STAR Methods

**Figure S2.** Single-cell RNA-seq cluster assignments, Related to Figure 1

**Figure S3.** Comparison of single-cell multiome with public datasets, Related to Figures 1 and 2

**Figure S4.** Comparison of single-cell ATAC-seq data from cell subpopulations, Related to Figure 2

**Figure S5.** Comparison of single-cell RNA-seq data from eyes with and without dry AMD, Related to Figure 1

**Figure S6.** Transcription factor binding motif analysis in human RPE and choroid cell types, Related to Figure 2

**Figure S7.** Enhancer mapping quality control metrics, Related to Figures 3 and 4

**Figure S8.** STARR-seq quality control metrics, Related to Figure 6

**Figure S9.** Characterization of STARR-seq enhancers, Related to Figure 6

**Figure S10.** ChromBPNet models of RPE and choroid cell types, Related to Figure 7

**Table S1.** Donor information, Related to STAR Methods

**Table S2.** Cell counts per sample, Related to Figure 1

**Table S3.** STARR-seq primers, Related to STAR Methods

**Protocol S1.** Isolation of single nuclei from frozen RPE-choroid, Related to STAR Methods

**Data S1.** Marker genes from scRNA-seq by cell type, Related to Figure 1

**Data S2.** Bed format of scATAC-seq peaks by cell type, Related to Figure 2

**Data S3.** Differentially expressed genes in macular RPE compared to peripheral RPE, Related to Figure S2

**Data S4.** Differentially expressed genes in AMD compared to control cell types, Related to Figure S5

**Data S5.** Bed format of scATAC-seq peaks co-accessible with promoter, Related to Figure 2

**Data S6.** Bed format of scATAC-seq peaks correlated with expression, Related to Figure 2

**Data S7.** −log10 adjusted p-values for TF motifs by cell type, Related to Figure S6

**Data S8.** H3K27ac HiChIP loops and intersecting genes, Related to Figure 3

**Data S9.** Enhancer-gene connections predicted by Activity-by-Contact, Related to Figure 3

**Data S10.** Prioritization of AMD risk variants, Related to Figures 5-7

**Data S11.** Allele-specific STARR-seq activity, Related to Figure 6

