## Supplemental Information for "Single-cell multiome and enhancer connectome of human retinal pigment epithelium and choroid nominate pathogenic variants in age-related macular degeneration"

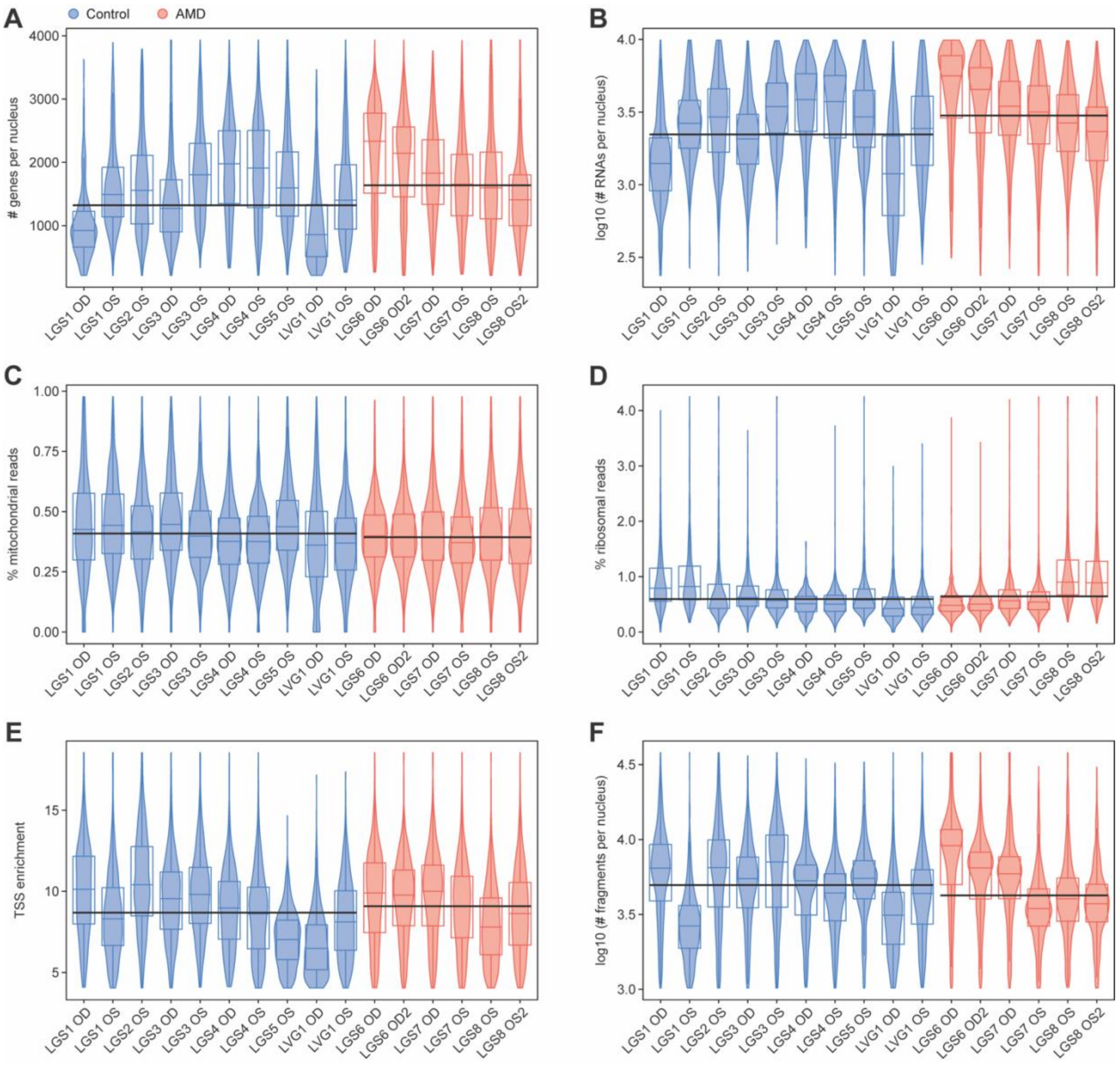


**Figure S1. Single-cell RNA- and ATAC-seq quality control metrics, Related to STAR Methods.**

**(A-F)** Violin plots depicting the number of detected genes per nucleus (A), number of detected RNA transcripts per nucleus (B), percent of mitochondrial reads (C), percent of ribosomal reads (D), TSS enrichment (E), and number of detected fragments per nucleus (F) by sample and disease status in the final dataset. Boxes depict the 25th percentile, median, and 75th percentile of each metric. Black lines depict the median of each metric across control or AMD samples. For two eyes (LGS6OD and LGS8OS), a second set of scRNA- and scATAC-seq libraries were generated to increase the number of nuclei profiled.

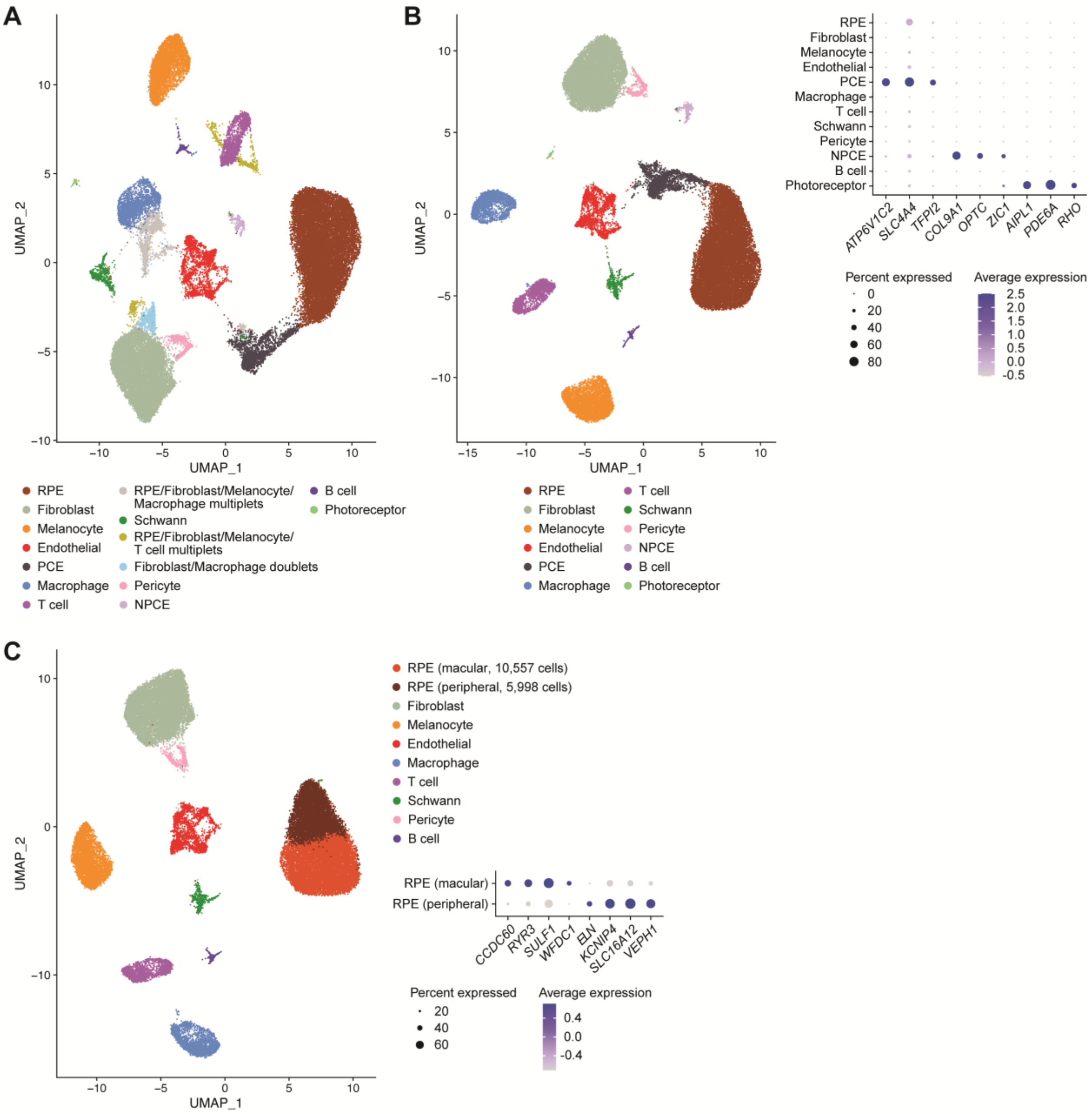


**Figure S2. Single-cell RNA-seq cluster assignments, Related to Figure 1.**

**(A)** First iteration uniform manifold approximation and projection (UMAP) plot of the scRNA-seq dataset after quality control filtering, automated doublet removal, and exclusion of clusters with no detected marker genes. Three clusters comprised of putative doublets or multiplets were removed and the dataset re-clustered. **(B)** Second iteration UMAP of the scRNA-seq dataset. Three clusters of non-RPE-choroid cell types (pigmented ciliary epithelium [PCE], nonpigmented ciliary epithelium [NPCE], and photoreceptor) were identified and removed and the dataset re-clustered. Dot plot depicts the normalized RNA expression by cluster of non-RPE-choroid marker genes.[1–3] The color and size of each dot signify the average expression level and percent of expressing cells, respectively. **(C)** Final iteration UMAP of the scRNA-seq dataset. Two clusters of RPE cells representing macular and peripheral subpopulations were identified and grouped together for downstream analyses. Dot plot depicts the normalized RNA expression of macular and peripheral RPE marker genes in RPE subpopulations.[4,5]

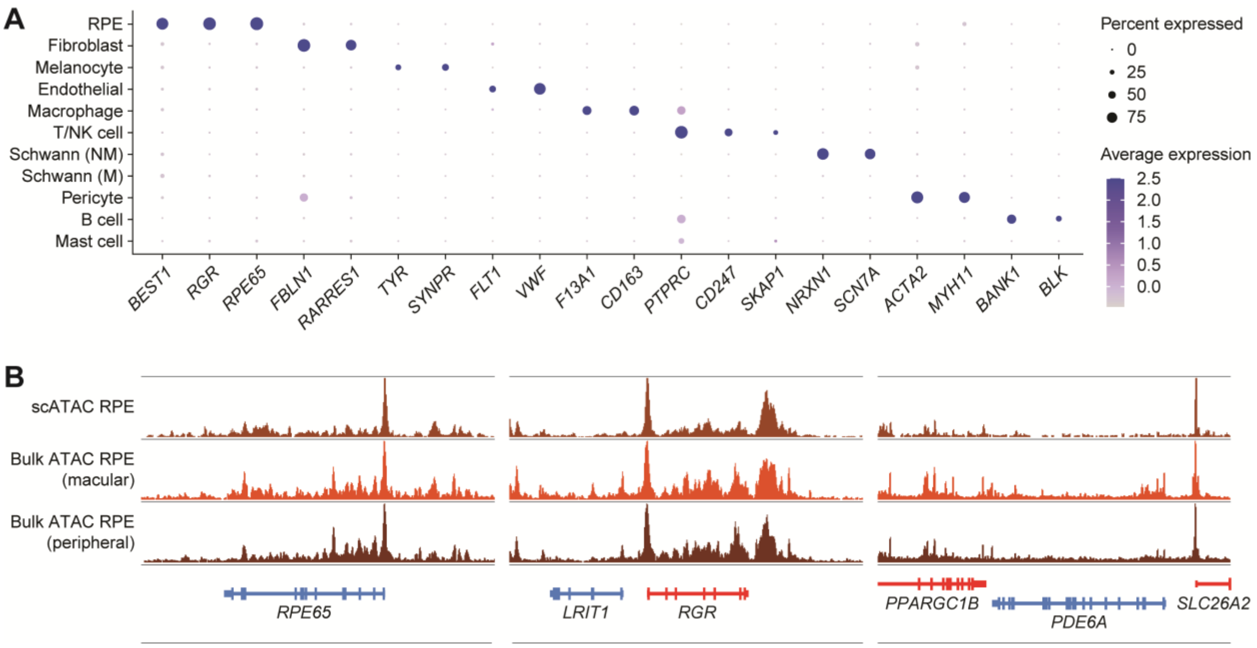


**Figure S3. Comparison of single-cell multiome with public datasets, Related to Figures 1 and 2.**

**(A)** Dot plot depicting the normalized RNA expression of the same marker genes from Figure 1D in a scRNA-seq dataset of human RPE and choroid from Voigt et al.[6] NK, natural killer; NM, non-myelinated; M, myelinated. **(B)** Sequencing tracks of chromatin accessibility comparing scATAC-seq of RPE from this study with bulk ATAC-seq of macular (GSM2640487) and peripheral (GSM2640485) human RPE from Wang et al.[7] Genes in the sense and antisense directions are shown in red and blue, respectively. Coordinates for each region: *RPE65* (chr1:68417805-68464772), *RGR* (chr10:84225860-84275121), *PDE6A* (chr5:149800089-149978610).

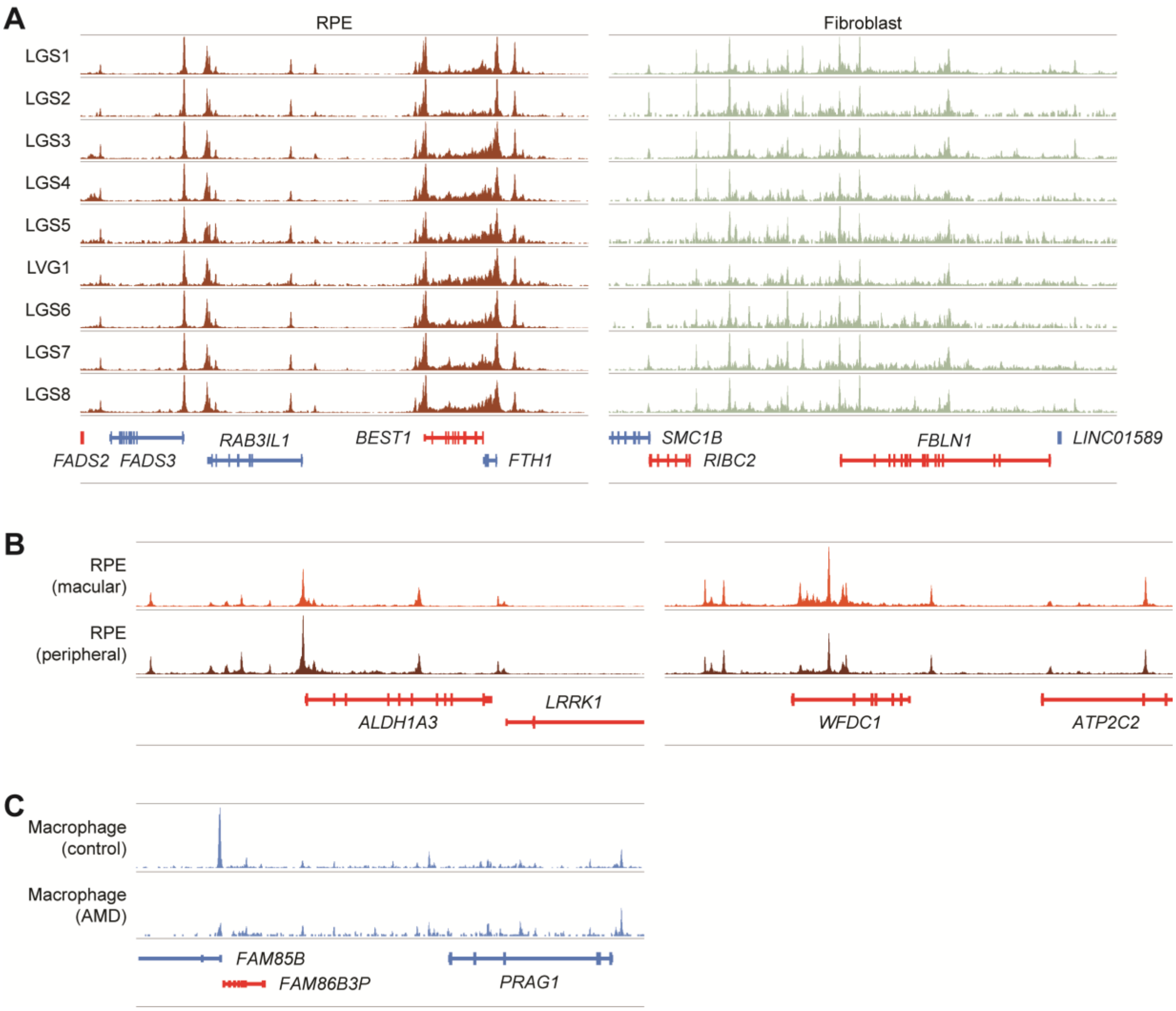


**Figure S4. Comparison of single-cell ATAC-seq data from cell subpopulations, Related to Figure 2.**

**(A)** Sequencing tracks of cell type chromatin accessibility by donor. Genes in the sense and antisense directions are shown in red and blue, respectively. Tracks were generated for cell types with >200 cells per donor. Coordinates for each region: RPE (chr11:61866483-61989817), fibroblast (chr22:45395132-45632169). **(B)** Sequencing tracks comparing chromatin accessibility in macular RPE cells versus peripheral RPE cells. Macular and peripheral RPE subpopulations were assigned by expression of marker genes shown in Figure S2C. Coordinates for each region: left (chr15:100846962-100946962), right (chr16:84254774-84404774). **(C)** Sequencing tracks comparing chromatin accessibility in choroidal macrophages from control versus AMD donors. Coordinates for region: chr8:8195477-8395477.

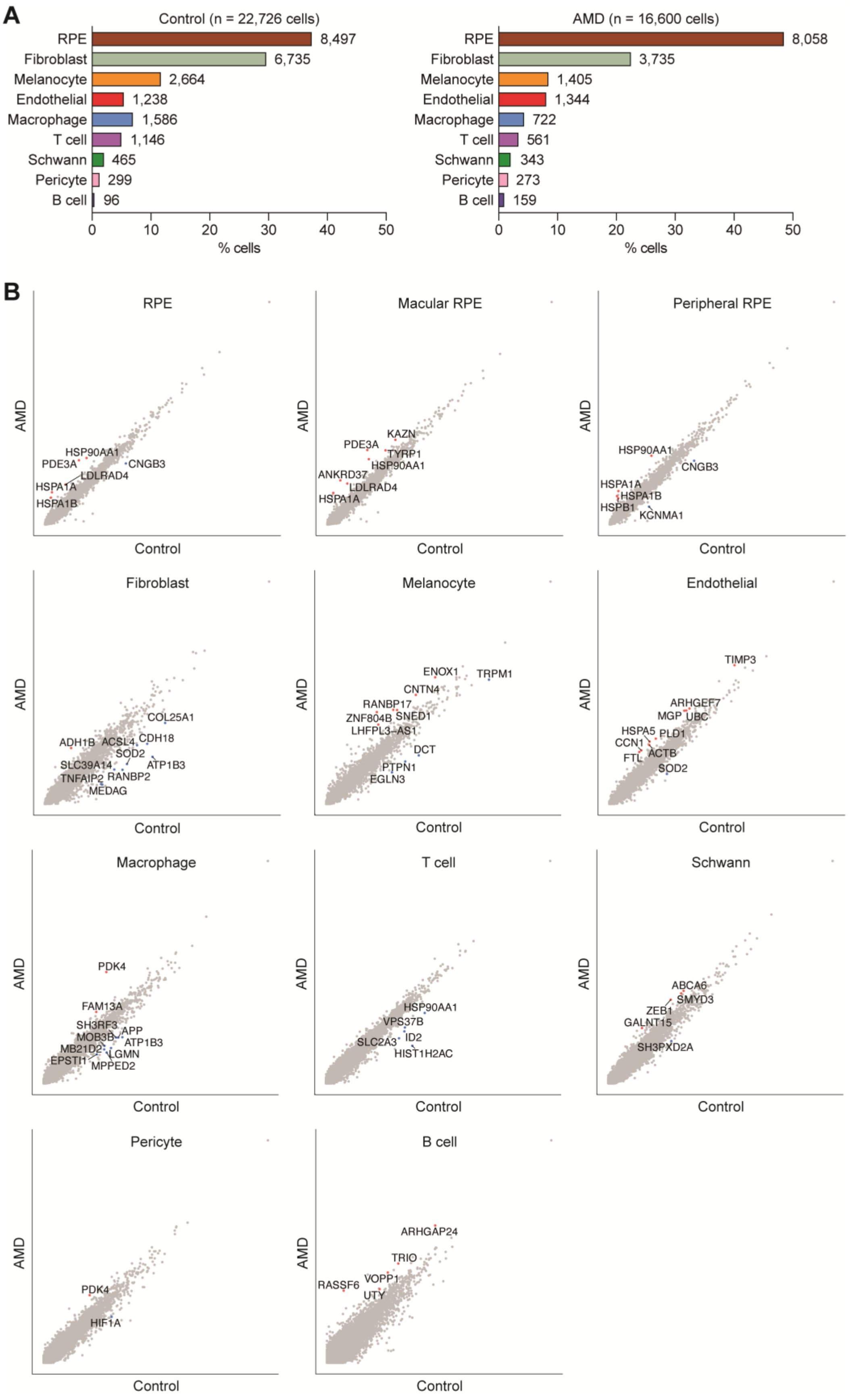
A
Control (n = 22,726 cells)
RPE 8,497
Fibroblast 6,735
Melanocyte 2,664
Endothelial 1,238
Macrophage 1,586
T cell 1,146
Schwann 465
Pericyte 299
B cell 96
% cells
AMD (n = 16,600 cells)
RPE 8,058
Fibroblast 3,735
Melanocyte 1,405
Endothelial 1,344
Macrophage 722
T cell 561
Schwann 343
Pericyte 273
B cell 159
% cells
B
RPE
Macular RPE
Peripheral RPE
Fibroblast
Melanocyte
Endothelial
Macrophage
T cell
Schwann
Pericyte
B cell
AMD
Control

**Figure S5. Comparison of single-cell RNA-seq data from eyes with and without dry AMD, Related to Figure 1.**

**(A)** Frequency of different human RPE and choroid cell types from control versus dry AMD eyes. Numbers adjacent to each bar indicate absolute counts. **(B)** Scatter plots comparing the average expression of each gene by cell type in control versus AMD eyes. Each dot represents a gene. Genes differentially expressed (log2 fold change ≥0.5, minimum fraction 0.3) in control and AMD cells are labeled in blue and orange, respectively. Up to ten genes with the highest absolute fold change for each cell type are shown.

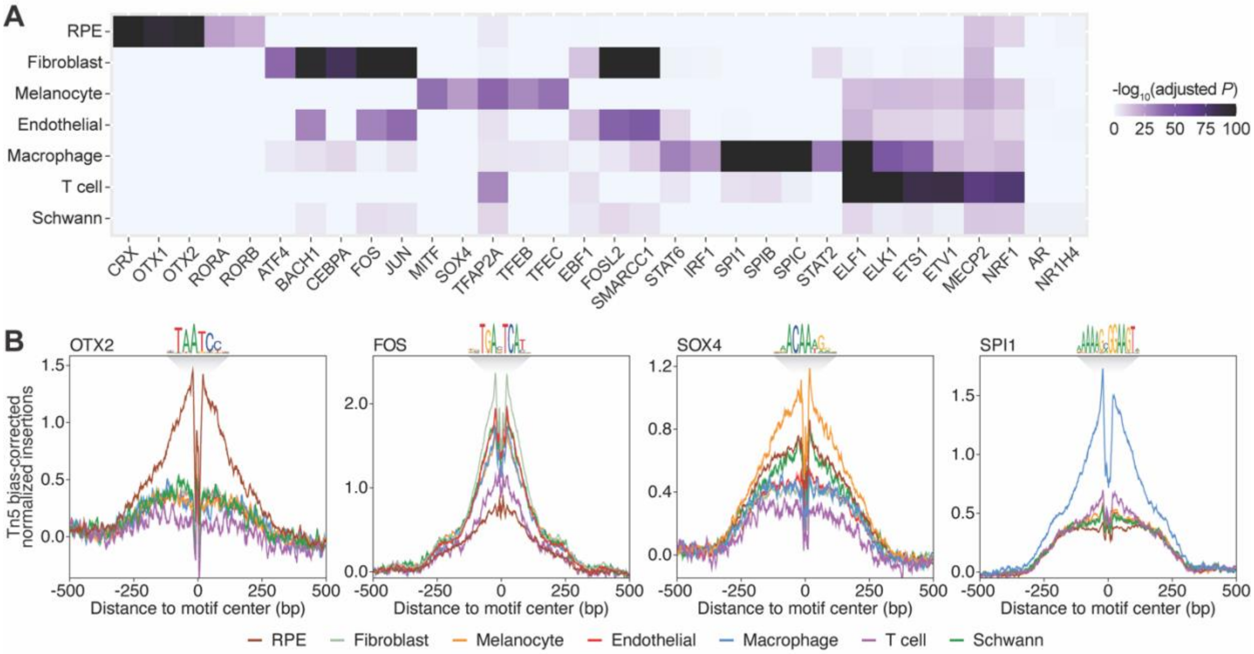


**Figure S6. Transcription factor binding motif analysis in human RPE and choroid cell types, Related to Figure 2**

**(A)** Heatmap of selected transcription factor binding motifs enriched in scATAC-seq peaks from RPE and choroid cell types. Darker colors indicate more significant enrichment. **(B)** Footprinting analysis of selected transcription factors across RPE and choroid cell types. Footprints were corrected for Tn5 insertion bias by subtracting the Tn5 insertion signal from the footprinting signal.

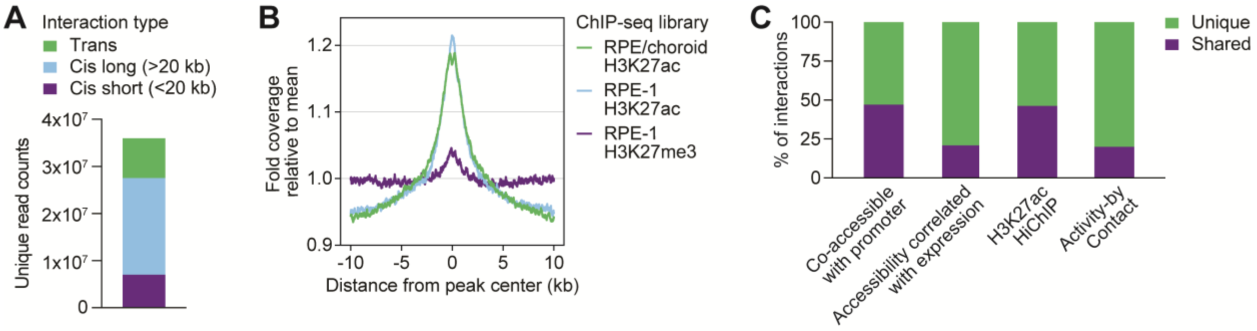


**Figure S7. Enhancer mapping quality control metrics, Related to Figures 3 and 4**

**(A)** Number and characterization of interactions identified by H3K27ac HiChIP. Cis refers to interactions on the same chromosome, while trans refers to interactions spanning separate chromosomes. **(B)** H3K27ac HiChIP read coverage around peak centers from published ChIP-seq of human RPE and choroid tissue or the RPE-1 hTERT cell line.[8,9] Aggregated read coverage at each base position was normalized to the mean coverage within 10 kb of peak centers. **(C)** Overlap of enhancer mapping strategies. Shared was defined as interactions from different methods having overlapping enhancers and the same target gene. Analysis of H3K27ac HiChIP was confined to loops containing at least one gene.

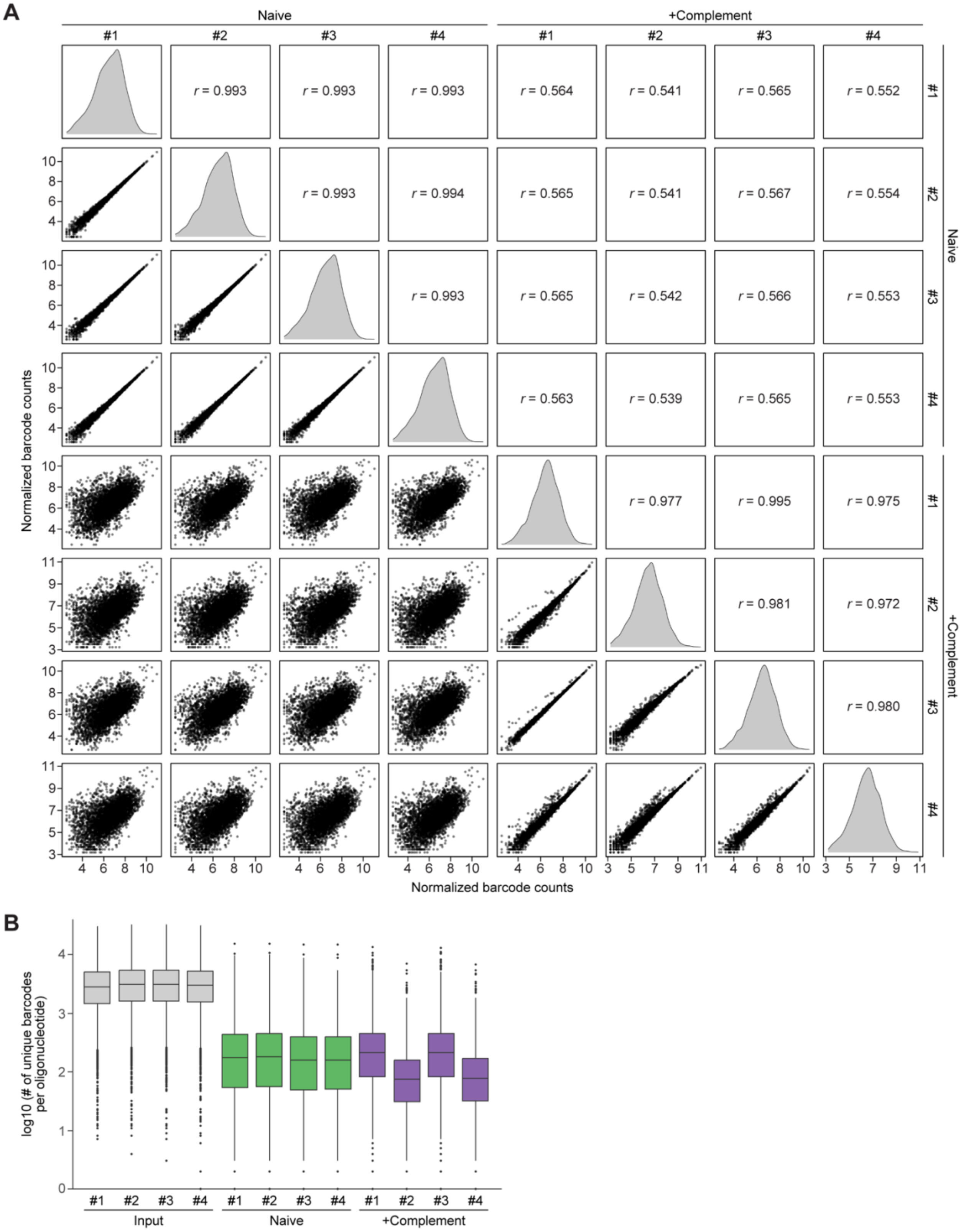


**Figure S8. STARR-seq quality control metrics, Related to Figure 6**

**(A)** Pairwise correlations of normalized barcode counts among STARR-seq output libraries. Histograms of normalized counts are shown in gray. *r* values indicate Pearson correlation coefficients. **(B)** Number of unique barcodes per oligonucleotide across STARR-seq input and output libraries. Boxes depict the 25th percentile, median, and 75th percentile.

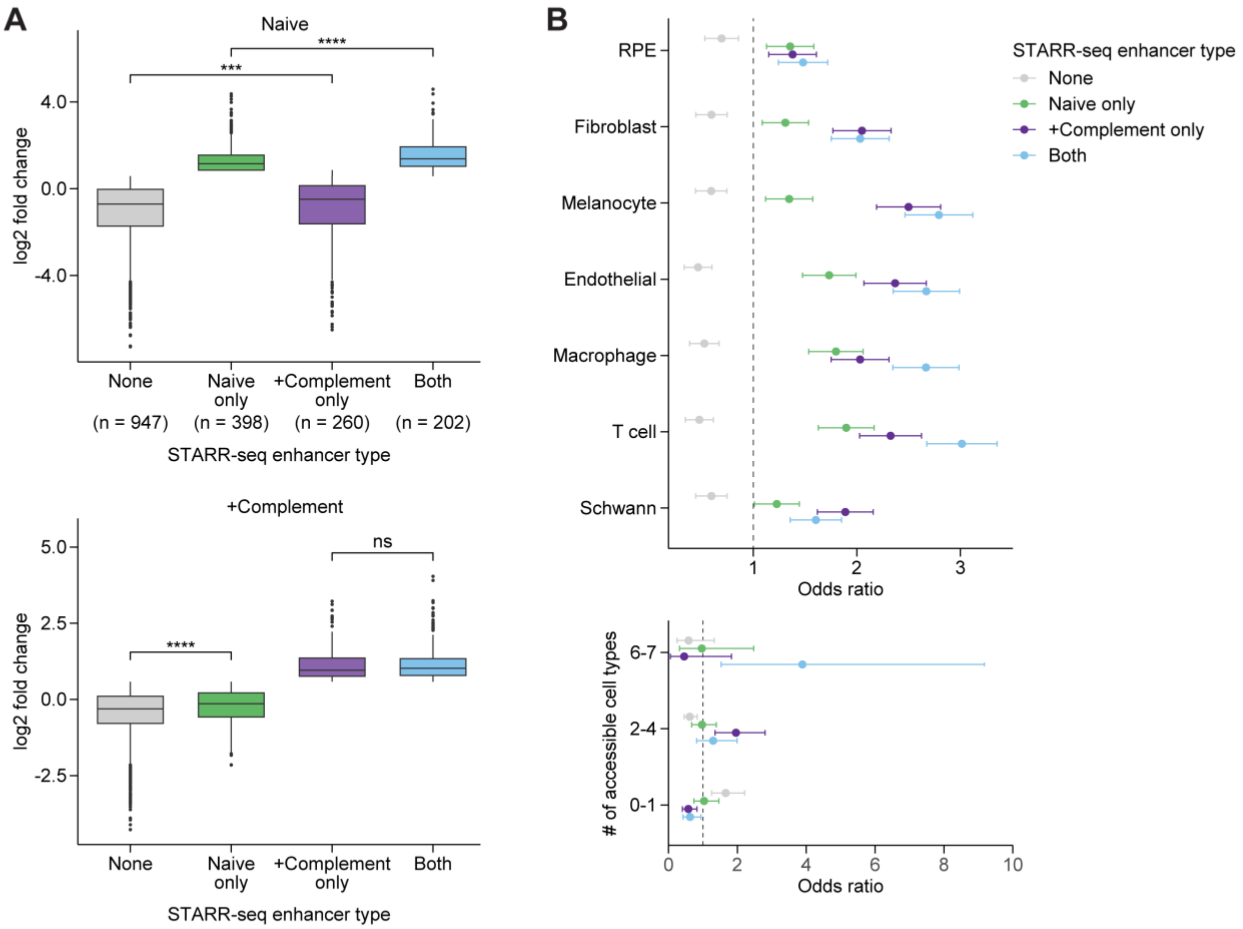


**Figure S9. Characterization of STARR-seq enhancers, Related to Figure 6**

**(A)** Regulatory activity of all oligonucleotides passing quality control filters in naive (top) and complement-activated (bottom) RPE cells by STARR-seq enhancer type. STARR-seq enhancers were defined as oligonucleotides exhibiting an absolute fold change >1.5 (log2 fold change >0.585) and adjusted p-value <0.05 in output libraries relative to the input library. Enhancers were grouped by whether they met this definition in only naive RPE cells, only complement-activated RPE cells, or both conditions. Boxes depict the 25th percentile, median, and 75th percentile. *** p <0.001, **** p <0.0001 by Wilcoxon rank-sum test with adjustment for multiple comparisons. ns, not significant. **(B)** Enrichment of STARR-seq enhancers for RPE and choroid scATAC-seq peaks. Bars depict 95% confidence intervals.

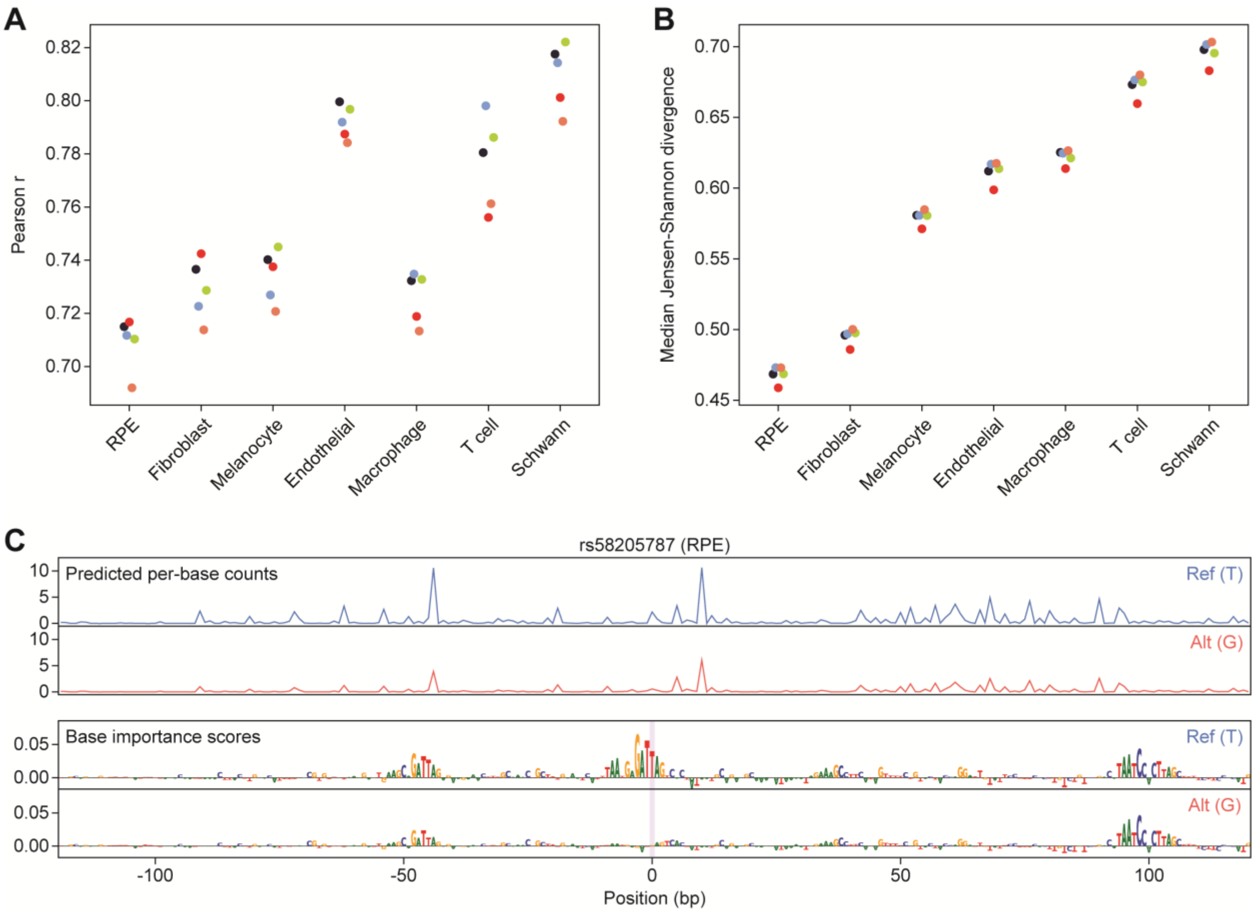


**Figure S10. ChromBPNet models of RPE and choroid cell types, Related to Figure 7**

**(A,B)** ChromBPNet performance metrics. Correlation scores between predicted and observed log counts (A) and Jensen-Shannon divergence scores between predicted and observed profiles (B) in peak regions that were withheld during cell type-specific model training. Each color represents one of five model folds. Higher scores for (A) and lower scores for (B) indicate more accurate model performance. **(C)** High effect variant example. Top: predicted per-base accessibility for reference (blue) and alternative (orange) alleles of rs58205787 (chr7:105075043) in RPE as determined by ChromBPNet models. Bottom: depiction of the importance of each base to predicted accessibility for reference and alternative alleles. The location of rs58205787 is highlighted in purple.

### Table S1. Donor information, Related to STAR Methods

| ID | Age | Sex | Death-to-preservation | Cause of death | Dry AMD? | Stage |
|---|---|---|---|---|---|---|
| LGS1 | 85 | F | 11 hours | Sepsis | No | -- |
| LGS2 | 87 | M | 9 hours | Fall | No | -- |
| LGS3 | 74 | F | 12 hours | Dementia | No | -- |
| LGS4 | 69 | M | 6 hours | Respiratory failure | No | -- |
| LGS5 | 89 | M | 9 hours | Sepsis | No | -- |
| LVG1 | 55 | M | 9 hours | Cardiac arrest | No | -- |
| LGS6 | 95 | M | 12 hours | Dementia | Yes | Early or intermediate |
| LGS7 | 91 | M | 16 hours | Dementia | Yes | Early or intermediate |
| LGS8 | 76 | F | 14 hours | Multi-organ failure | Yes | Advanced |

**Table S2. Cell counts per sample, Related to Figure 1**

| | LGS1 OD | LGS1 OS | LGS2 OS | LGS3 OD | LGS3 OS | LGS4 OD | LGS4 OS | LGS5 OS |
|---|---|---|---|---|---|---|---|---|
| RPE | 1498 | 586 | 620 | 980 | 331 | 964 | 734 | 1103 |
| Fibroblast | 1679 | 711 | 240 | 878 | 321 | 476 | 417 | 578 |
| Melanocyte | 629 | 122 | 150 | 578 | 127 | 219 | 183 | 251 |
| Endothelial | 289 | 52 | 104 | 374 | 77 | 84 | 68 | 78 |
| Macrophage | 468 | 133 | 91 | 137 | 72 | 125 | 159 | 132 |
| T cell | 211 | 90 | 154 | 65 | 72 | 68 | 132 | 62 |
| Schwann | 95 | 21 | 27 | 83 | 6 | 66 | 13 | 87 |
| Pericyte | 50 | 20 | 13 | 53 | 14 | 47 | 28 | 21 |
| B cell | 23 | 8 | 9 | 15 | 2 | 3 | 7 | 6 |
| Total | 4942 | 1743 | 1408 | 3163 | 1022 | 2052 | 1741 | 2318 |

| | LVG1 OD | LVG1 OS | LGS6 OD | LGS6 OD2 | LGS7 OD | LGS7 OS | LGS8 OS | LGS8 OS2 |
|---|---|---|---|---|---|---|---|---|
| RPE | 911 | 770 | 375 | 1883 | 1083 | 2058 | 889 | 1770 |
| Fibroblast | 868 | 567 | 69 | 323 | 417 | 1060 | 519 | 1347 |
| Melanocyte | 201 | 204 | 58 | 211 | 178 | 325 | 127 | 506 |
| Endothelial | 30 | 82 | 29 | 156 | 85 | 224 | 319 | 531 |
| Macrophage | 133 | 136 | 21 | 22 | 98 | 201 | 100 | 280 |
| T cell | 109 | 183 | 27 | 148 | 59 | 149 | 63 | 115 |
| Schwann | 41 | 26 | 10 | 41 | 52 | 86 | 62 | 92 |
| Pericyte | 33 | 20 | 14 | 74 | 25 | 42 | 27 | 91 |
| B cell | 11 | 12 | 8 | 41 | 21 | 64 | 11 | 14 |
| Total | 2337 | 2000 | 611 | 2899 | 2018 | 4209 | 2117 | 4746 |

**Table S3. STARR-seq primers, Related to STAR Methods**

| Name | Sequence |
|---|---|
| STARR-seq-1F | ATTAGATTGATCTAGAGCATGCACCGGTacatttctctatcgataggtac |
| STARR-seq-1R | TCGAAGCGGCCGGCCGAATTCGTCGA[barcode]CTCTACACGCAATGGCTAGC |
| RT | CAAACTCATCAATGTATCTTATCATG |
| STARR-seq-2F | ACACTCTTTCCCTACACGACGCTCTTCCGATCTattagattgatctagagcatgcaccggt |
| STARR-seq-2R | GTGACTGGAGTTCAGACGTGTGCTCTTCCGATCtcgaagcggccggccgaattcgtcga |
| TruSeq universal | AATGATACGGCGACCACCGAGATCTACACTCTTTCCCTACACGACGCTCTTCCGATCT |
| Indexing | CAAGCAGAAGACGGCATACGAGAT[index]GTGACTGGAGTTCAGACGTG |

**Protocol S1. Isolation of single nuclei from frozen RPE-choroid, Related to STAR Methods.**
Adapted from dx.doi.org/10.17504/protocols.io.6t8herw

***Before you start the protocol:***

1. All steps should be performed on ice or at 4°C. Pre-chill a swinging bucket centrifuge and a fixed angle centrifuge to 4°C.
2. Pre-chill all Dounces and pestles to 4°C. For each sample, you will need three 2 mL glass Dounces, at least one "A" loose pestle, and at least one "B" tight pestle (Sigma-Aldrich D8938).
3. Pre-chill all tubes. For each sample, you will need one 25 or 50 mL conical tube for filtration and two 2 mL round-bottom LoBind tubes for gradient separation.
4. Prepare all buffers. For faster dissolution, crush protease inhibitor tablets prior to addition to 1x Homogenization Buffer Unstable Solution. DTT, spermidine, spermine, and digitonin are stored at -20°C. All other detergents, ATAC-RSB, and other buffers are stored at 4°C.
5. Fill a beaker with sterile water to soak used Dounces and pestles.

***Isolation of Nuclei via Dounce Homogenization and Density Gradient Centrifugation:***

6. Remove samples from liquid nitrogen storage and keep on dry ice until use. Pre-chill a petri dish, a pair of forceps, and a razor blade on dry ice.
7. Add 1 mL Homogenization Buffer Unstable Solution (HBUS) to each glass Dounce on ice.
8. For each sample, transfer tissue onto the petri dish on dry ice and cut into rice grain-sized pieces with the pre-chilled razor blade. Using pre-chilled forceps, transfer pieces into glass Dounces. Aim for three rice grain-sized pieces per Dounce. Also transfer any pulverized tissue from petri dish to Dounces.
9. Add an additional 400 uL HBUS to each Dounce. Let tissues thaw in Dounces for 5 minutes.
10. Dounce with "A" loose pestle until resistance goes away (~10-20 strokes per Dounce). Place "A" pestle(s) into beaker with sterile water to soak for cleaning.
11. Dounce with "B" tight pestle for 20 strokes per Dounce. Place "B" pestle(s) into beaker with sterile water to soak for cleaning.
12. Filter homogenate through a 70 um cell strainer into a 25 or 50 mL conical tube pre-chilled on ice.
13. Transfer filtrate to two 2 mL LoBind tubes pre-chilled on ice.
14. Pellet nuclei by spinning 5 minutes at 4°C at 350 RCF in a fixed angle centrifuge.
15. For each tube, remove all but ~50 uL supernatant, leaving nuclei pellet intact.
16. Gently resuspend nuclei in a total volume of 400 uL HBUS. If you left 50 uL in the previous step, this means you should add 350 uL HBUS. Fully resuspended without clumps using a P1000 pipette.
17. For each tube, add 1 volume (400 uL) 50% Iodixanol Solution and mix well by pipetting.
18. For each tube, slowly layer 600 uL 30% Iodixanol solution A under the 25% mixture. To avoid mixing layers, first wipe side of the pipette tip with a Kimwipe to remove excess.
19. For each tube, layer 600 uL 40% Iodixanol solution A under the 30% mixture. To avoid mixing layers, first wipe side of the pipette tip with a Kimwipe to remove excess. During this step, you will need to gradually draw your pipette tip up to avoid overflowing the tube. However, the tip of your pipette must stay below the 30%-40% interface at all times.
20. In a pre-chilled swinging bucket centrifuge, spin for 20 minutes at 4°C at 3,000 RCF with brakes off. Handle tubes gently so as to not disturb the gradient.

21. During spin, prepare Resuspension Buffer with 0.1% Tween and 2% BSA (RSB-TB) and pre-chill one 2 mL LoBind on ice.
22. Carefully remove tubes from the centrifuge and use a P1000 pipette to aspirate top layers of the gradient down to within 200-300 uL of the faint nuclei band at the 30%-40% interface. Be careful not to disrupt the nuclei band.
23. Using a P200 pipette, collect the faint nuclei band and transfer to a pre-chilled 2 mL LoBind tube containing 1.6 mL RSB-TB. Minimize transfer of pigment found immediately below the nuclei band and do not aspirate more than 200 uL per tube. Transfer nuclei bands from the same tissue sample to the same LoBind tube.
24. Pellet nuclei by spinning 5 minutes at 4°C at 350 RCF in a fixed angle centrifuge.
25. Remove all but ~50 uL supernatant, leaving nuclei pellet intact.
26. Gently resuspend nuclei in a total volume of 400 uL RSB-TB. Fully resuspended without clumps using a P1000 pipette.
27. Add 1 volume (400 uL) 50% Iodixanol Solution and mix well by pipetting.
28. Slowly layer 600 uL 30% Iodixanol solution B under the 25% mixture. Avoid mixing layers.
29. Layer 600 uL 40% Iodixanol solution B under the 30% mixture. Avoid mixing layers and gradually draw your pipette tip up to avoid overflowing the tube.
30. In a pre-chilled swinging bucket centrifuge, spin for 20 minutes at 4°C at 3,000 RCF with brakes off. Handle tube gently so as to not disturb the gradient.
31. During spin, pre-chill a 1.5 mL LoBind and a 5 mL polystyrene tube with 35 um strainer on ice (Falcon 352235). If using nuclei for single-nuclei ATAC (including single-nuclei multiome), prepare 0.1x Lysis Buffer and Diluted Nuclei Buffer.
32. Carefully remove tubes from the centrifuge and use a P1000 pipette to aspirate top layers of the gradient down to within 200-300 uL of the faint nuclei band at the 30%-40% interface. Be careful not to disrupt the nuclei band.
33. Using a P200 pipette, collect the faint nuclei band and transfer to a pre-chilled 1.5 mL LoBind tube containing 1.1 mL RSB-TB. Minimize transfer of pigment found immediately below the nuclei band and do not aspirate more than 200 uL.
34. Filter nuclei through the 35 um strainer into the pre-chilled polystyrene tube. Transfer filtrate back into the 1.5 mL LoBind tube.
35. Pellet nuclei by spinning 5 minutes at 4°C in a fixed angle centrifuge. Use 350 RCF if isolating bulk nuclei and 250 RCF if preparing a single-nuclei suspension.
36. Remove supernatant. Using a P1000 pipette, remove all but ~100 uL supernatant. Remove remaining supernatant with a P200 pipette set to 200 uL using a single fluid pipetting motion. Place pipette tip on the side of tube opposite to the nuclei pellet during the P200 aspiration.
37. If isolating bulk nuclei, the nuclei pellet can be processed immediately or flash-frozen on dry ice and stored at -80°C.

***Using Nuclei for Single-Nuclei ATAC or Multiome with 10x Genomics:***

38. Resuspend pellet in 200 uL 0.1x Lysis Buffer and pipette mix 5 times. Incubate on ice for 2 minutes.
39. During incubation, pre-chill a 5 mL polystyrene tube with 10 um strainer on ice (Fisher Scientific NC1721506). Pre-wet strainer with 200 uL RSB-TB.
40. Wash nuclei with 900 uL RSB-TB and pipette mix 5 times. Filter nuclei through the 10 um strainer into the pre-chilled polystyrene tube. Transfer filtrate back into the 1.5 mL LoBind tube.

41. Wash the polystyrene tube with an additional 200 uL RSB-TB and transfer to the 1.5 mL LoBind tube.
42. Pellet nuclei by spinning 5 minutes at 4°C at 250 RCF in a fixed angle centrifuge. Pellet will be very small but should be visible.
43. Remove supernatant. Using a P1000 pipette, remove all but ~100 uL supernatant. Remove remaining supernatant with a P200 pipette set to 200 uL using a single fluid pipetting motion. Place pipette tip on the side of tube opposite to the nuclei pellet during the P200 aspiration.
44. Resuspend pellet in at least 12 uL of Diluted Nuclei Buffer depending on size of the pellet. Gently pipette mix until pellet completely dissolves.
45. Combine 5 uL of nuclei suspension with 5 uL of Trypan blue in a separate tube. Load mixture into a disposable manual hemocytometer and count nuclei. An ideal preparation will have few nuclei aggregates.
46. If needed, reduce nuclei concentration by adding additional Diluted Nuclei Buffer. We recommend using 5 uL at ~1,610 nuclei/uL for a targeted nuclei recovery of 5,000.
47. Proceed with 10x Genomics protocol.

***Stock Buffers:***
Recipes for stock buffers (1.034x Homogenization Buffer Stable Solution, Diluent Buffer, 50% Iodixanol Solution, and ATAC-RSB Buffer) can be found at:
https://www.protocols.io/view/isolation-of-nuclei-from-frozen-tissue-for-atac-se-kxygxmr34l8j/v1/materials

***Same Day Buffers:***
Homogenization Buffer Unstable Solution (HBUS) – 25 mL

| *Stock* | *Name* | *Volume (uL)* | *Catalog* |
|---|---|---|---|
| 1.034x | Homogenization Buffer Stable Solution | 23,925 | - |
| 1 M | DTT | 25 | Thermo Scientific R0861 |
| 500 mM | spermidine | 25 | Sigma S2501 |
| 150 mM | spermine | 25 | Sigma S3256 |
| 10% | NP40 (IGEPAL CA-630) | 750 | Sigma I8896 |
| - | cOmplete Protease Inhibitor | 0.5 tablet | Roche 11697498001 |
| - | RNase Inhibitor (replace with Homogenization Buffer Stable Solution if omitting RNA) | 250 | New England Biolabs M0314 |

30% Iodixanol Solution A – 1.5 mL

| *Stock* | *Name* | *Volume (uL)* | *Catalog* |
|---|---|---|---|
| - | HBUS | 600 | - |
| 50% | Iodixanol Solution | 900 | - |

40% Iodixanol Solution A – 1.5 mL

| *Stock* | *Name* | *Volume (uL)* | *Catalog* |
|---|---|---|---|
| - | HBUS | 300 | - |
| 50% | Iodixanol Solution | 1,200 | - |

Resuspension Buffer with 0.1% Tween and 2% BSA (RSB-TB) – 8 mL

| *Stock* | *Name* | *Volume (uL)* | *Catalog* |
|---|---|---|---|
| - | ATAC-RSB | 6,240 | - |
| 10% | Tween-20 | 80 | Roche 11332465001 |
| 10% | MACS BSA Stock Solution | 1,600 | Miltenyi Biotec 130-091-376 |
| - | RNase Inhibitor (replace with ATAC-RSB if omitting RNA) | 80 | New England Biolabs M0314 |

30% Iodixanol Solution B – 750 uL

| *Stock* | *Name* | *Volume (uL)* | *Catalog* |
|---|---|---|---|
| - | RSB-TB | 300 | - |
| 50% | Iodixanol Solution | 450 | - |

40% Iodixanol Solution B – 750 uL

| *Stock* | *Name* | *Volume (uL)* | *Catalog* |
|---|---|---|---|
| - | RSB-TB | 150 | - |
| 50% | Iodixanol Solution | 600 | - |

Recipes for 0.1x Lysis Buffer and Diluted Nuclei Buffer can be found at: https://cdn.10xgenomics.com/image/upload/v1660261285/support-documents/CG000375_DemonstratedProtocol_NucleiIsolationComplexSample_ATAC_GEX_Sequencing_Rev_C.pdf
